# Functional characterization of terpene synthases from *Dendrobium nobile* provides insights into the biosynthesis of copaborneol, a proposed dendrobine precursor

**DOI:** 10.64898/2026.10.06.757176

**Authors:** Dandan Li, Jianling Wang, Sijing Fu, Hualei Wang

## Abstract

Picrotoxane-type sesquiterpene alkaloids, including dendrobine, are characteristic constituents of *Dendrobium nobile*, but the enzymes that produce their proposed skeletal precursors remain incompletely characterized. Here, we combined tissue metabolite profiling with genome-guided screening and biochemical characterization of terpene synthases (TPSs). GC-MS analysis revealed variation in sesquiterpenoid profiles and dendrobine signals among tissues and developmental stages. Manual curation of the chromosome-level genome yielded 16 full-length TPS-a candidates. Functional screening in an FPP-supplying *Escherichia coli* system showed that three closely related genes on chromosome 1, *DnTPS1*–*DnTPS3*, produced copaborneol as their dominant product. The principal product of *DnTPS1* was purified and identified by NMR spectroscopy, and a purified-enzyme assay confirmed its direct conversion of FPP to copaborneol. Fluorescence imaging in rice protoplasts and *Nicotiana benthamiana* leaves indicated predominantly cytosolic localization of DnTPS1. Guided by sequence comparisons and structural modeling, we tested 12 DnTPS1 variants and found that substitutions at W377 and I295 altered the relative distribution of sesquiterpene products. Phylogenetic and regional gene-order analyses placed DnTPS1∼DnTPS3 in the same clade and identified related TPS candidates within corresponding genomic regions of six Dendrobium species. These findings establish that three *D. nobile* TPSs can form copaborneol and identify DnTPS1 residues that influence product selectivity, providing a basis for testing the proposed connection to dendrobine biosynthesis.

## 1. Introduction

Picrotoxane natural products are structurally diverse sesquiterpenoids that include non-nitrogenous members such as tutin and alkaloids such as dendrobine (Mou et al., 2021; Shi et al., 2022). Dendrobine is a characteristic constituent of *Dendrobium nobile* and has attracted interest because of its reported biological activities (Bhardwaj et al., 2024; Li et al., 2022; Mou et al., 2021; Shi et al., 2023; Shi et al., 2025). However, the enzymes that generate its proposed terpenoid precursors remain incompletely characterized.

Classical isotope-feeding studies established a terpenoid origin for tutin and dendrobine (Arigoni, 1975; Corbella et al., 1969; Corbella et al., 1972; Corbella et al., 1973; Corbella et al., 1975; Edwards et al., 1970; Turnbull et al., 1972; Yamazaki et al., 1966). Labeled copaborneol was converted to tutin in *Coriaria japonica* (Turnbull et al., 1972), but the analogous conversion to dendrobine has not been demonstrated in *D. nobile*. The FPP-derived cyclization step and the subsequent oxidative and nitrogen-incorporating reactions therefore need to be evaluated with direct enzyme and in planta evidence.

Plant terpene synthases (TPSs) convert farnesyl diphosphate (FPP) into diverse sesquiterpenes through metal-ion-dependent ionization, cyclization, rearrangement and reaction termination (Christianson, 2017; Degenhardt et al., 2009; Karunanithi and Zerbe, 2019). The conserved DDXXD and NSE/DTE motifs initiate substrate ionization, whereas the active-site cavity governs subsequent carbocation rearrangements and product formation. Thus, identifying the TPS responsible for copaborneol formation is an important step toward understanding dendrobine biosynthesis. Copaborneol has been reported among the products of TPSs from maize and Medicago (Garms et al., 2010; Saldivar et al., 2023). Comparisons of closely related *D. nobile* TPSs and tests of residues affecting their product distribution can therefore extend the biochemical understanding of copaborneol formation.

In this study, we screened 16 manually curated, full-length terpene synthase candidates from the chromosome-level *D. nobile* genome by heterologous expression and GC-MS analysis. Three closely related genes on chromosome 1, *DnTPS1*∼*DnTPS3*, produced copaborneol as their major detectable product. The principal product of *DnTPS1* was purified and identified by NMR spectroscopy, and its conversion of FPP to copaborneol was confirmed with purified enzyme. Transient expression in *N. benthamiana* supported copaborneol formation in a plant system, while fluorescence assays in rice protoplasts and *N. benthamiana* leaves indicated predominantly cytosolic localization. Guided by sequence comparisons and structural modeling, we tested 12 DnTPS1 variants and found that substitutions at W377 and I295 altered the relative distribution of its sesquiterpene products. Together, these results identify enzymes capable of producing copaborneol in *D. nobile* and provide a basis for investigating whether this compound contributes to dendrobine biosynthesis.

## 2. Results

### 2.1 Tissue-specific profiles of sesquiterpenoids and dendrobine in Dendrobium nobile

To examine tissue-and developmental-stage variation in sesquiterpenoid profiles, seeds, protocorms, tissue-cultured seedlings, one-year-old stems, young leaves, and flowers of *D. nobile* were analyzed by headspace solid-phase microextraction coupled with gas chromatography-mass spectrometry (HS-SPME-GC-MS). A total of 27 compounds were tentatively annotated across the six sample types, comprising 26 sesquiterpenoids and dendrobine (compound 27) (Fig. 1, Table S1). Based on the annotated peaks, flowers exhibited the greatest diversity of detected sesquiterpenoids (11 compounds), followed by seeds (10), tissue-cultured seedlings (8), and protocorms and stems (7 each). Only one annotated sesquiterpenoid was detected in leaves.

**Fig. 1.**
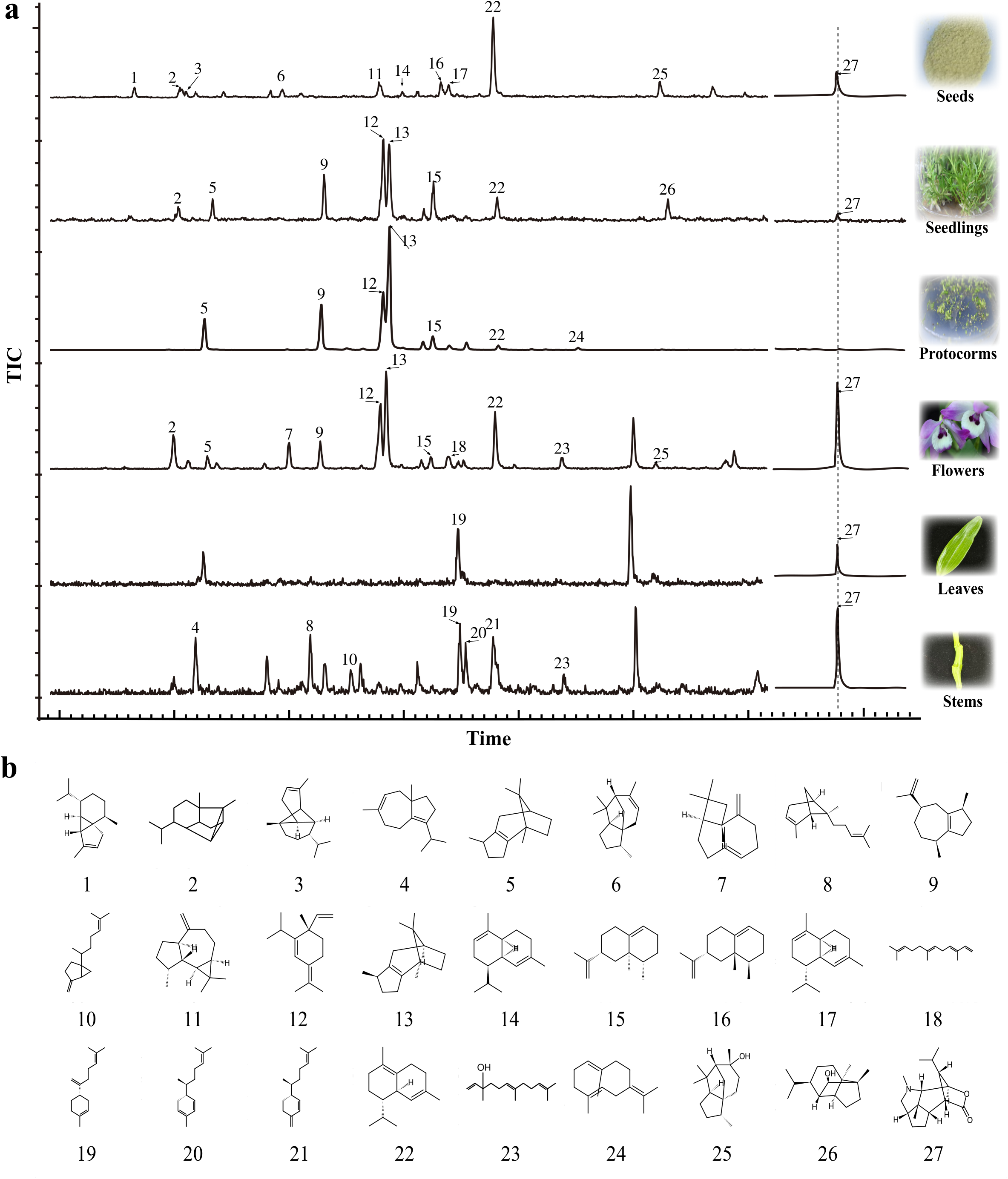
Overview of sesquiterpenoids and dendrobine in different tissues and developmental stages of *Dendrobium nobile*. (a) Representative HS-SPME-GC-MS total ion chromatograms of seeds, tissue-cultured plantlets, protocorms, flowers, leaves, and one-year-old stems. Blue traces show the volatile components, whereas the black trace on the right highlights the peak position of dendrobine. Numbers correspond to the compound structures shown in panel b. (b) Chemical structures of the putative sesquiterpenoids numbered 1∼24 and dendrobine described in the text. Peak assignments are tentative unless supported by authentic standards or additional structural evidence.

The distribution of individual compounds differed among the sample types. Compounds 1, 3, 6, 11, 14, 16, and 17 were detected only in seeds, whereas compound 26 was detected only in tissue-cultured seedlings and compound 24 only in protocorms. Compounds 7 and 18 were restricted to flowers, while compounds 4, 8, 10, 20, and 21 were detected only in stems. Compounds 5, 9, 12, 13, 15, and 22 were shared by tissue-cultured seedlings, protocorms, and flowers; compound 22 was also detected in seeds. Compound 2 occurred in seeds, tissue-cultured seedlings, and flowers, whereas compound 25 was detected in seeds and flowers. Compound 23 was shared by flowers and stems, and compound 19 was detected in both leaves and stems. The peak tentatively assigned to dendrobine (compound 27) was observed in all sample types except protocorms, with a comparatively weak signal in tissue-cultured seedlings. The corresponding electron-ionization mass spectra are provided in Supplementary Fig. S1. These profiles indicate tissue-and stage-associated differences in the occurrence of tentatively annotated sesquiterpenoids and dendrobine in *D. nobile*.

### 2.2 Identification and sequence characterization of full-length TPS genes of D. nobile

To identify candidate enzymes for sesquiterpene formation, we examined 33 putative TPS-a genes annotated in the chromosome-level *D. nobile* genome (Li et al., 2024). Assisted by GSAman, manual curation yielded 16 full-length candidate sequences with complete open reading frames and the expected class I TPS domains. Phylogenetic analysis separated the 16 DnTPS proteins into three clusters (Fig. 2a). DnTPS1, DnTPS2, and DnTPS3 formed a distinct cluster and were located within the same chromosome 1 region. All 16 proteins retained the DDXXD and NSE/DTE metal-binding motifs, although their broader motif patterns varied (Fig. 2b). In particular, DnTPS1–DnTPS3 contained motif 14 and lacked motif 10, whereas DnTPS44 lacked both motifs; the remaining DnTPS proteins contained motif 10 but lacked motif 14. Alignment of regions associated with metal binding also showed a residue pattern shared by DnTPS1∼DnTPS3 (Fig. 2c).

**Fig. 2.**
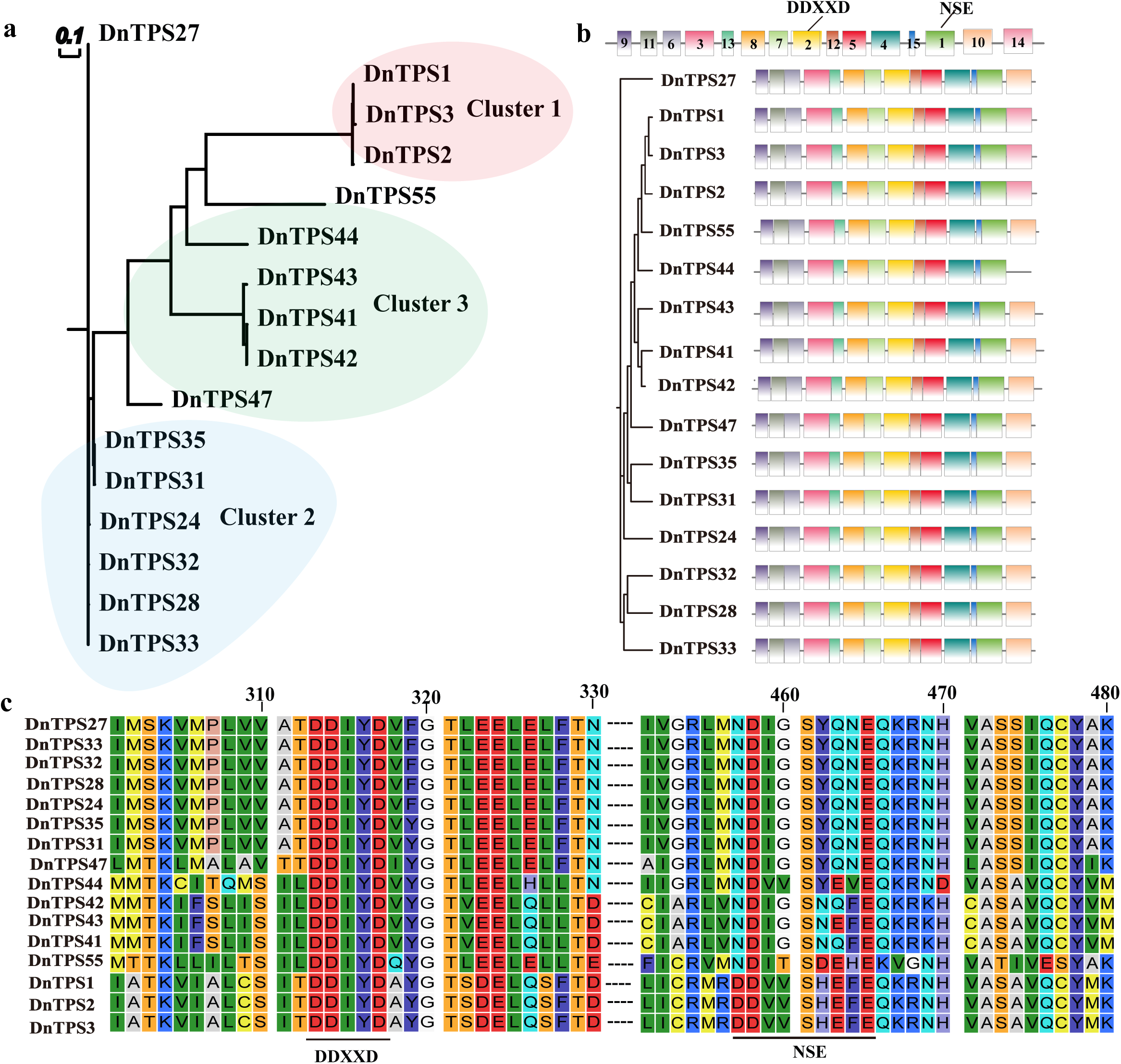
Sequence characteristics of 16 manually curated full-length TPS candidate proteins from *Dendrobium nobile*. (a) Phylogenetic relationships among the candidate proteins. The scale bar represents the number of substitutions per site. (b) Distribution of conserved motifs. Different colored blocks represent MEME motifs with distinct motif numbers, and the positions of the DDXXD and NSE/DTE motifs are indicated. (c) Amino acid sequence alignment of regions associated with metal ion binding. DnTPS1–DnTPS3 differ from the other candidates at several positions.

### 2.3. Functional screening identifies three D. nobile copaborneol synthases

Functional screening of the 16 curated *DnTPS* candidates in the FPP-supplying *E. coli* C41 system showed that *DnTPS1*–*DnTPS3* generated copaborneol as their dominant detectable product (Fig. 3a). *DnTPS1* also produced peaks #1 and #2. Six chromosome 10 candidates yielded peak #4, provisionally assigned as daucene (Fig. 3b). On chromosome 12, *DnTPS42* generated peaks #5∼#8, whereas *DnTPS44* generated peak #9 (Fig. 3c). Proposed structures corresponding to peaks #1∼#9 are shown in Fig. 3d, and their EI mass spectra are provided in Supplementary Fig. S2. No target sesquiterpene product was detected for *DnTPS41*, *DnTPS43* or *DnTPS47* under the screening conditions.

**Fig. 3.**
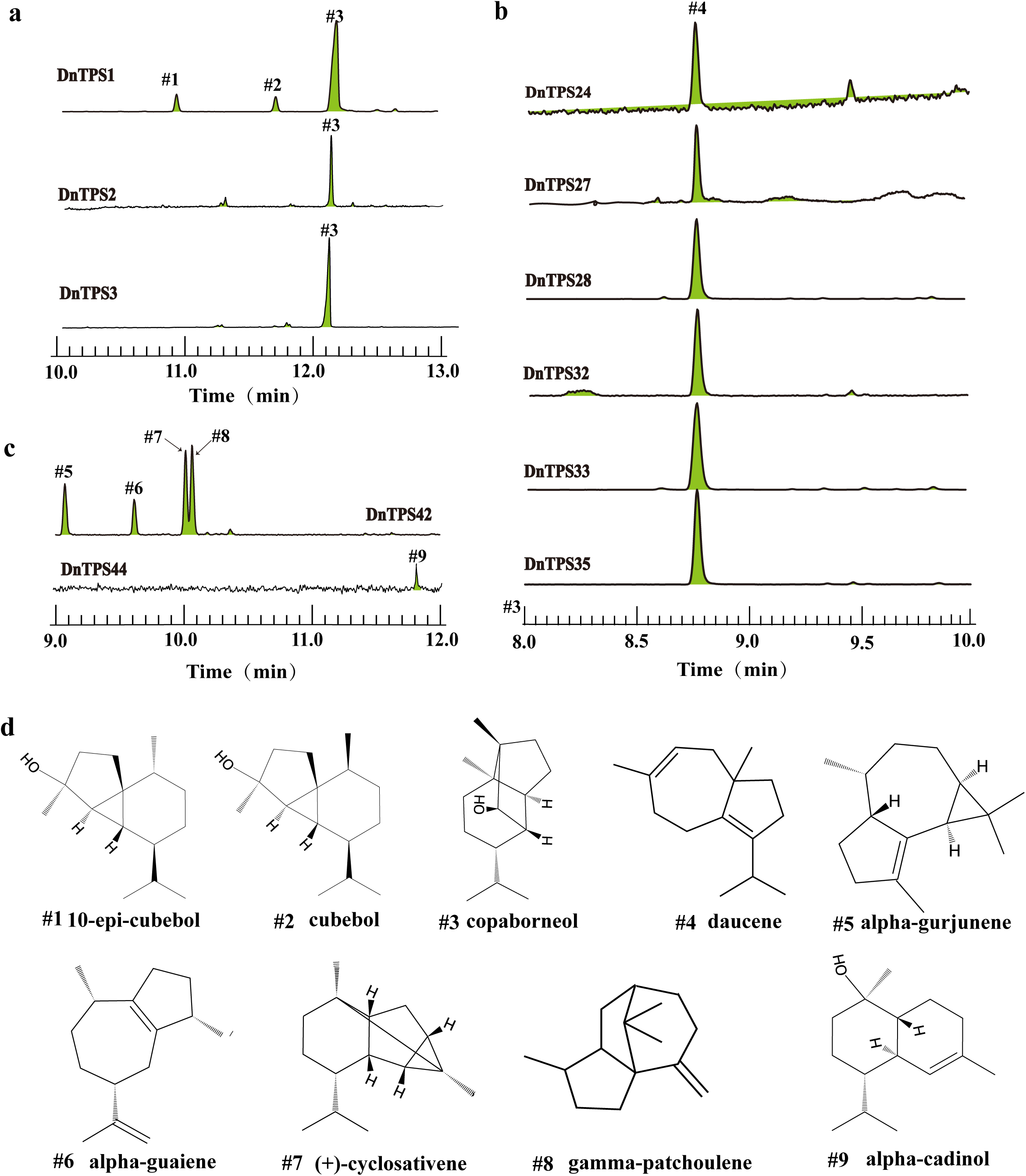
Heterologous expression product profiles of *TPS* candidate genes from *Dendrobium nobile*. (a) Products formed by DnTPS1–DnTPS3 in an FPP-supplying *Escherichia coli* system. (b) Product profiles of six candidate *TPS* genes located on chromosome 10. (c) Product profiles of *DnTPS42* and *DnTPS44*. (d) Proposed chemical structures and names corresponding to peaks #1∼#9; the major DnTPS1 product (#3) was further supported by NMR.

Because DnTPS1∼DnTPS3 share approximately higher than 95% protein-sequence identity and have the same dominant detectable product, DnTPS1 was chosen for further characterization. Its major product was purified and assigned as copaborneol by comparison of its ^1^H and ^13^C NMR spectra with published data (Lusi et al., 2022) (Supplementary Figs. S3 and S4). The dominant products of DnTPS2 and DnTPS3 were assigned as copaborneol on the basis of matching GC-MS retention behavior and mass spectra relative to the structurally characterized DnTPS1 product.

### 2.4. Purified DnTPS1 catalyzes copaborneol formation and localizes to the cytosol

To verify copaborneol formation independently of the whole-cell screening system, recombinant DnTPS1 was expressed in *Escherichia coli BL21* and purified. In the presence of FPP, purified DnTPS1 generated a product matching copaborneol by chromatographic analysis, supporting direct catalysis of the conversion of FPP to copaborneol (Fig. 4a). We further expressed *DnTPS1* in *Nicotiana benthamiana*, where copaborneol production was detected (Fig. 4b). In protoplasts isolated from etiolated rice seedlings, the DnTPS1-GFP fluorescence signal was predominantly detected in the cytoplasm (Fig. 4c). Subcellular localization analysis was also performed in *N. benthamiana* leaves (Fig. 4d), and the fluorescence pattern was likewise consistent with a predominantly cytosolic distribution and with the predictions of TargetP and WoLF PSORT (Supplementary Table S2). Together, these purified-enzyme, heterologous expression, and subcellular localization experiments support DnTPS1 as a copaborneol-forming enzyme with predominantly cytosolic fluorescence in the tested heterologous plant cells.

**Fig. 4.**
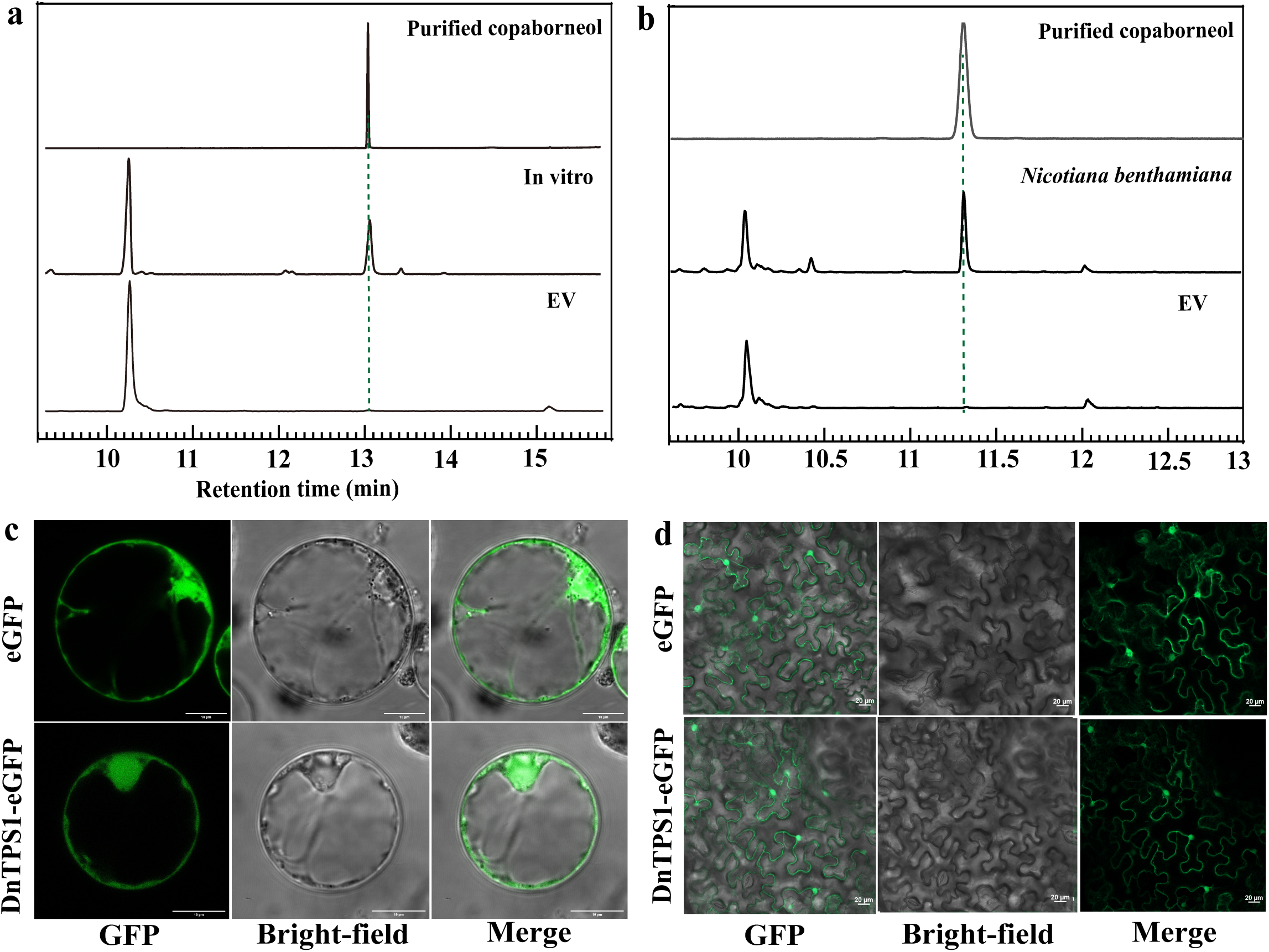
In vitro catalysis, transient expression in *Nicotiana benthamiana*, and subcellular localization of DnTPS1. (a) Chromatographic comparison of purified copaborneol, the in vitro reaction of purified DnTPS1 with FPP, and the empty-vector control; the dashed line indicates the retention time corresponding to copaborneol. (b) Detection of products from transient expression in *N. benthamiana*, compared with purified copaborneol and the empty-vector control. (c) Fluorescence, bright-field, and merged images of free eGFP and DnTPS1-eGFP in protoplasts isolated from etiolated rice seedlings. (d) Fluorescence imaging of the same constructs in *N. benthamiana* leaves. Scale bars are as indicated in the images.

### 2.5. Identification of key residues affecting the product selectivity of DnTPS1

After establishing that DnTPS1 catalyzes the conversion of FPP to copaborneol, we further combined sequence comparison, structural modeling, and site-directed mutagenesis to investigate amino acid residues potentially involved in determining its product selectivity. Multiple sequence alignment showed that DnTPS1 and related homologs from Dendrobium species shared similar residues at several positions, whereas these positions differed from the corresponding residues in other functionally characterized sesquiterpene synthases (Fig. 5a). Based on the spatial distribution of residues in the DnTPS1–FPP complex model, Y12, I295, C298, W377, T402, T403, and C404 were selected for site-directed mutagenesis (Fig. 5b). A total of 12 mutants were generated, and their product profiles were compared by heterologous expression followed by GC-MS analysis (Fig. 5c,d).

**Fig. 5.**
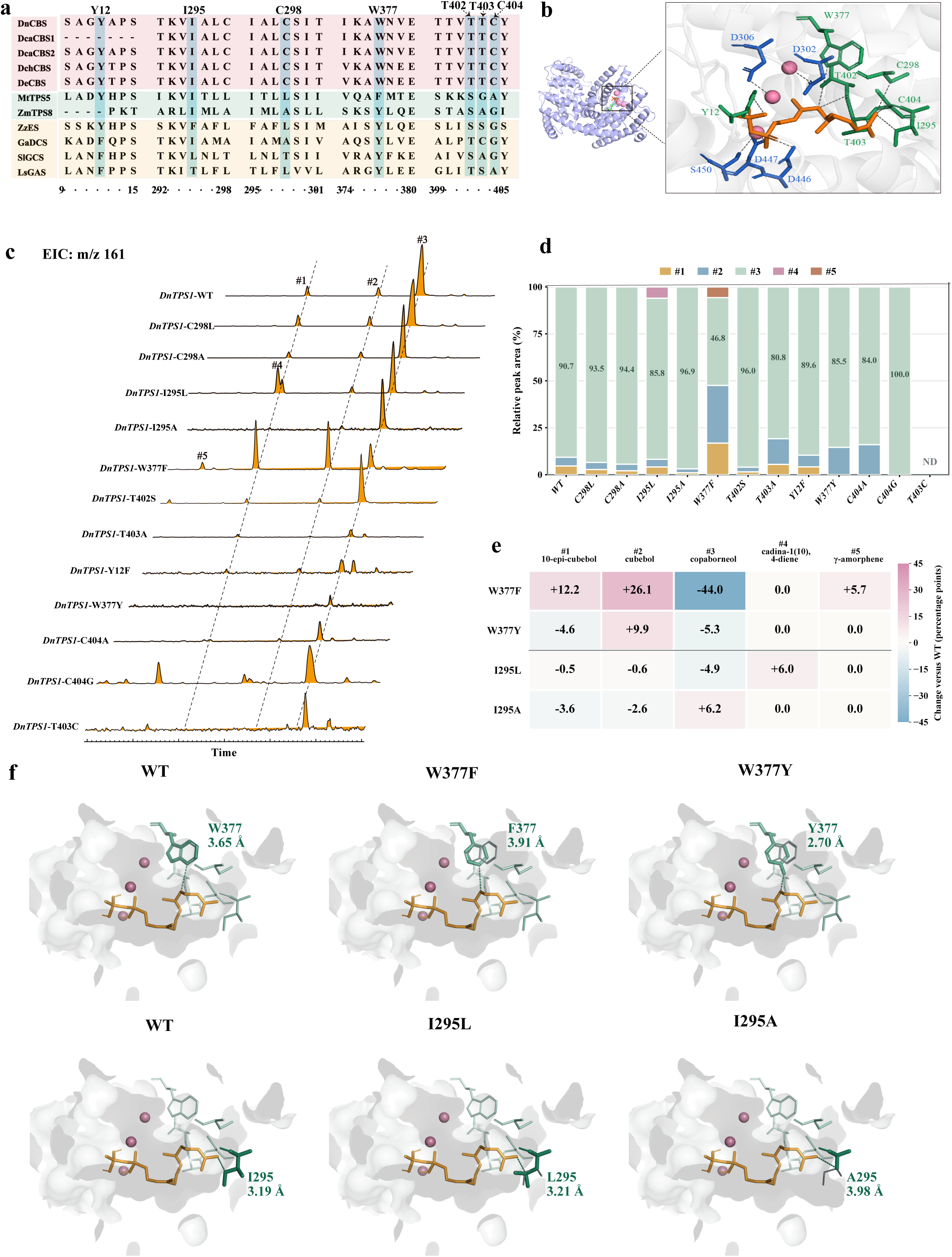
Effects of amino acid substitutions on the product selectivity of DnTPS1. (a) Partial amino acid sequence alignment of DnTPS1, related TPS candidates from Dendrobium species, and previously characterized sesquiterpene synthases. Arrows indicate Y12, I295, C298, W377, T402, T403, and C404 examined in this study. (b) Overall structure of the predicted DnTPS1-FPP complex and an enlarged view of the substrate-binding pocket. FPP is shown as orange sticks, candidate residues as green sticks, acidic residues of the DDXXD and NSE/DTE motifs as blue sticks, and metal ions as pink spheres. (c) Extracted-ion chromatograms (EICs; m/z 161) of products generated by wild-type DnTPS1 (WT) and 12 amino acid variants. Peaks 1∼5 correspond to 10-epi-cubebol, cubebol, copaborneol, cadina-1(10),4-diene, and γ-amorphene, respectively. (d) Relative peak-area compositions of the five target products generated by WT and the corresponding variants. Product proportions were normalized to the summed peak area of the five target products. ND indicates that none of the target products was detected under the conditions used. (e) Changes in the relative proportions of individual products in W377F, W377Y, I295L, and I295A compared with WT. Values represent percentage-point differences. (f) Predicted local structures of the substrate-binding pockets of WT, W377F, W377Y, I295L, and I295A. The indicated values are predicted distances between FPP and the corresponding amino acid residue. The structural models illustrate potential local spatial changes associated with the substitutions.

Wild-type DnTPS1 predominantly produced copaborneol (#3), which accounted for 90.7% of the total peak area of the five analyzed products. Amino acid substitutions at different positions affected the product profile to varying degrees, with W377F showing the most pronounced change. Compared with the wild type, the relative peak area of copaborneol in W377F decreased by 44.0 percentage points, whereas those of 10-epi-cubebol (#1) and cubebol (#2) increased by 12.2 and 26.1 percentage points, respectively. In addition, γ-amorphene (#5) appeared as a new detectable product, accounting for 5.7% of the analyzed products. By contrast, copaborneol remained the predominant product of W377Y, although its relative peak area decreased by 5.3 percentage points. The proportion of cubebol increased by 9.9 percentage points, whereas that of 10-epi-cubebol decreased by 4.6 percentage points (Fig. 5c-e). These results indicate that different side-chain substitutions at W377 have distinct effects on product partitioning, with W377F causing a marked reduction in the relative abundance of copaborneol among the analyzed products.

The two substitutions at I295 also resulted in distinct changes in product composition. I295L reduced the relative peak area of copaborneol by 4.9 percentage points and led to the appearance of cadina-1(10),4-diene (#4), which accounted for 6.0% of the analyzed products. In contrast, I295A increased the relative peak area of copaborneol by 6.2 percentage points, accompanied by decreases of 3.6 and 2.6 percentage points in 10-epi-cubebol and cubebol, respectively (Fig. 5d,e). Among the remaining mutants, C298L, C298A, and T402S retained product profiles dominated by copaborneol. Y12F, T403A, W377Y, and C404A also produced copaborneol as the major product, although the relative proportions of minor products were altered. Among the five target products included in the analysis, only copaborneol was detected for C404G, whereas none of these target products was detected for T403C under the current analytical conditions (Fig. 5c,d).

To further examine the local structural changes associated with substitutions at W377 and I295, the substrate-binding pockets of the wild type and the corresponding mutants were compared. In the structural models, the indicated distance between W377 and FPP was 3.65 Å, whereas the corresponding distances for W377F and W377Y were 3.91 and 2.70 Å, respectively. For I295, I295L, and I295A, the corresponding distances were 3.19, 3.21, and 3.98 Å, respectively (Fig. 5f). These modeled differences suggest that the substitutions may alter the local spatial environment surrounding the substrate. Together with the mutant product profiles, these results identify W377 and I295 as important residues affecting the product selectivity of DnTPS1. However, their precise roles in controlling catalytic intermediates and individual reaction steps remain to be experimentally resolved.

### 2.6. Phylogenetic relationships and local genomic organization of DnTPS1∼DnTPS3-related loci across Dendrobium species

To compare the relationships of DnTPS1–DnTPS3 with homologous TPS proteins from other Dendrobium species, we constructed a maximum-likelihood phylogenetic tree using 15 proteins (≥300 aa) that were highly homologous to DnTPS1–DnTPS3 and located within the selected genomic regions of six species. The results showed that DnTPS1, DnTPS2, and DnTPS3 clustered within the same clade, with SH-aLRT/UFBoot support values of 94.2/100. This clade was adjacent to the clade containing the *Dendrobium huoshanense* candidates Dhu000014629 and Dhu000014627 (Fig. 6a).

**Fig. 6.**
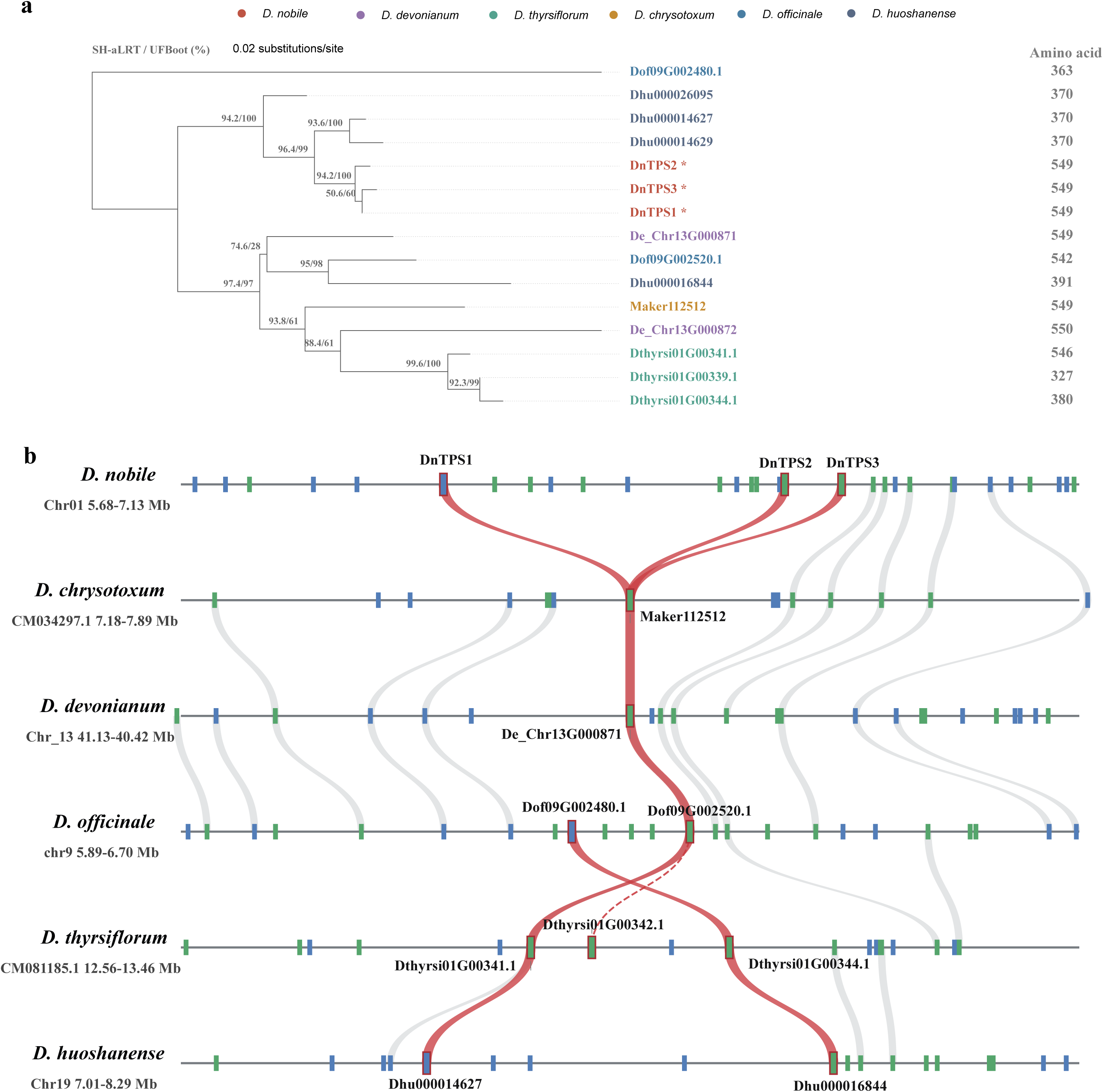
Phylogenetic relationships and local genomic organization of DnTPS1–DnTPS3-related loci across Dendrobium species. (a) Maximum-likelihood phylogeny of 15 TPS candidate proteins from selected genomic regions of six Dendrobium species. Only proteins of ≥300 aa were included. The MAFFT alignment was trimmed with ClipKIT to 410 amino acid sites, and the tree was inferred in IQ-TREE under the Q.plant+G4 model. Node labels indicate SH-aLRT/UFBoot support values based on 1,000 replicates each. The scale bar represents amino acid substitutions per site, and midpoint rooting was used for visualization. Tip colors indicate species, values on the right indicate protein lengths, and asterisks mark DnTPS1– DnTPS3, which were biochemically characterized in this study. (b) Local genomic organization of DnTPS1–DnTPS3-related regions in six Dendrobium species. Blue and green gene blocks indicate genes on the forward and reverse strands, respectively, and red outlines identify focal TPS candidates. Grey ribbons connect ordered regional reciprocal-best-hit anchors. Each species track is scaled independently; horizontal distances cannot be compared between species. Solid red ribbons indicate protein-homology matches among focal TPS candidates, including the one-to-many relationships between DnTPS1–DnTPS3 and Maker112512. The dashed red line indicates a partial protein match between Dthyrsi01G00342.1 and Dof09G002520.1, covering less than 50% of the longer protein.

Comparison of gene positions showed that *DnTPS1*–*DnTPS3* were all located within the selected region of chromosome 1 (Chr01) in *D. nobile*. Several background homologous genes with similar gene order were observed across the corresponding regions of the six species (Fig. 6b), consistent with local gene-order correspondence among these genomic regions. The red TPS links further showed that all three DnTPS proteins exhibited high protein sequence similarity to Maker112512 from *D. chrysotoxum*, with local alignment identities of 86.2%, 86.1%, and 86.0%, respectively. Additional local homologous matches were also observed between the three DnTPS proteins and the TPS candidates shown in the corresponding regions of the other species.

## 3. Discussion

Terpene synthases (TPSs) convert common isoprenoid precursors into diverse carbon skeletons and therefore represent a major enzymatic source of structural diversity in plant sesquiterpenes (Christianson, 2017; Degenhardt et al., 2009; Karunanithi and Zerbe, 2019). In this study, functional screening of 16 manually curated full-length TPS candidates from *Dendrobium nobile* identified DnTPS1, DnTPS2, and DnTPS3 as enzymes that predominantly produce copaborneol. The catalytic function of DnTPS1 was supported by multiple lines of evidence, including isolation and NMR characterization of its major product, in vitro assays using purified recombinant protein and FPP, and heterologous expression in *Nicotiana benthamiana*. The major products of DnTPS2 and DnTPS3 were assigned as copaborneol based on their GC-MS characteristics, which were indistinguishable from those of the DnTPS1 product. Thus, three highly similar homologous enzymes retain the same major product type, providing a useful system for examining both functional conservation and variation in product selectivity among *D. nobile* TPSs.

The ability to form copaborneol is not unique to Dendrobium. The previously characterized maize enzyme ZmTPS8 produces multiple sesquiterpenes, including copaborneol, and contributes to fungus-induced defense-related metabolism (Saldivar et al., 2023). The contribution of the present study lies in the functional characterization of copaborneol-forming TPSs by comparing 16 related *D. nobile* candidates and the experimental analysis of DnTPS1 product selectivity through site-directed mutagenesis. Under the analytical and normalization conditions used here, copaborneol accounted for 90.7% of the total peak area of the five quantified products generated by wild-type DnTPS1, indicating a strongly copaborneol-biased product profile. This relatively selective catalytic behavior also provided a suitable baseline for evaluating how individual amino acid substitutions alter product partitioning.

The possible connection between DnTPS1–DnTPS3 and dendrobine-associated metabolism provides an additional biological context for these enzymes. Earlier isotope-tracing experiments demonstrated that labeled copaborneol could be converted into tutin in *Coriaria japonica* (Turnbull et al., 1972), providing evidence that copaborneol can serve as an intermediate in the formation of picrotoxane-type sesquiterpenes. In the present study, the enzymatic link between FPP and copaborneol establishes an experimentally accessible entry point for investigating downstream modification reactions and for screening enzymes capable of further transforming copaborneol. The fluorescence pattern of DnTPS1 in heterologous plant cells was consistent with a predominantly cytosolic localization, and copaborneol was detectable after transient expression of DnTPS1 in *N. benthamiana*, indicating that the enzyme can generate its target product in a plant cellular environment. In parallel, the differences in sesquiterpene and dendrobine signals among tissues and developmental stages of *D. nobile* point to pronounced spatiotemporal heterogeneity in specialized metabolism. However, these observations do not by themselves establish a direct biosynthetic relationship between DnTPS1–DnTPS3 and dendrobine formation in planta, and such a relationship will require additional experimental validation.

Site-directed mutagenesis identified W377 as an important determinant of DnTPS1 product distribution. Replacement of W377 with phenylalanine decreased the relative proportion of copaborneol by 44.0 percentage points, while increasing the proportions of 10-epi-cubebol and cubebol and leading to the detectable formation of γ-amorphene. In contrast, W377Y retained copaborneol as the predominant product. Notably, both substitutions preserve an aromatic side chain but result in clearly different product profiles. This observation suggests that the role of W377 cannot be explained solely by the presence or absence of an aromatic ring. Instead, differences in side-chain geometry, local polarity, and interactions with neighboring residues may contribute to the control of catalytic selectivity. The active sites of terpene synthases influence product outcome by constraining substrate conformations and by shaping the local environment in which highly reactive carbocation intermediates are generated, stabilized, and redirected toward alternative reaction branches (Christianson, 2017; Degenhardt et al., 2009). This general framework provides a plausible basis for interpreting the effects of substitutions at W377.

Substitutions at I295 produced a different pattern of changes in product composition. I295L decreased the relative proportion of copaborneol and led to the appearance of cadina-1(10),4-diene, whereas I295A increased the copaborneol proportion by 6.2 percentage points. These contrasting effects indicate that even substitutions among nonpolar residues can alter product partitioning and that the relationship between side-chain size and catalytic selectivity is not necessarily straightforward. Together with the effects observed at W377, these findings establish DnTPS1 as a useful enzymatic system for investigating how subtle changes in the local active-site environment influence sesquiterpene product branching. Previous work on MtTPS5 used isotopically labeled substrates, reactions in deuterated water, and mutational analyses to distinguish among alternative product-forming routes ^(Garms et al., 2010)^, illustrating the type of independent mechanistic evidence required to resolve individual catalytic pathways. For DnTPS1, the modeled changes in residue–substrate distances provide testable structural hypotheses, but they should not be regarded as direct evidence for specific interactions or reaction steps. Further mechanistic analyses will therefore be required to determine how W377 and I295 influence substrate conformation, carbocation rearrangement, or termination of the catalytic cascade.

Phylogenetic and local genomic-context analyses provide an evolutionary framework for these functional observations. DnTPS1∼DnTPS3 formed a well-supported clade in the phylogeny of TPS candidates from the selected genomic regions, and several ordered regional homolog matches were identified across the corresponding regions of the examined Dendrobium species. Together with the shared major product of DnTPS1∼DnTPS3 under the screening conditions, these observations provide a basis for functional comparisons with homologous TPS candidates from other Dendrobium species. At the same time, the changes in product distribution caused by individual amino acid substitutions illustrate how limited sequence divergence within a conserved TPS framework can potentially generate catalytic diversification.

## 4. Conclusions

In summary, this study functionally identified three *D. nobile* TPSs capable of producing copaborneol and demonstrated that substitutions at W377 and I295 are associated with altered product selectivity of DnTPS1. These enzymes and their engineered variants provide experimentally tractable materials for investigating the catalytic basis of copaborneol formation and sesquiterpene product diversification. In addition, establishing an enzymatic source of copaborneol provides a foundation for testing its potential role as a substrate for downstream reactions relevant to dendrobine-associated metabolism.

## 5. Experimental

### 5.1. Plant materials and sample collection

Seeds, protocorms, tissue-cultured seedlings, one-year-old young stems, young leaves, and flowers of *D. nobile* were collected in the plant growth chamber. Samples were harvested and immediately frozen in liquid nitrogen for RNA extraction. Similar fresh samples were harvested for SPME-GC-MS detection.

### 5.2. Profiling of volatile sesquiterpenoids

Metabolites were profiled for different tissues of *D. nobile* using headspace solid-phase microextraction coupled with gas chromatography-mass spectrometry (HS-SPME-GC-MS). 1.5 g fresh samples were placed in sealed headspace vials and pre-equilibrated at 60 °C for 10 min. An SPME fiber was then exposed to the headspace for 45 min. Following extraction, the fiber was immediately transferred to the GC inlet for thermal desorption at 250 °C for 3 min. Chromatographic separation was performed using an Agilent 6890 gas chromatograph equipped with an HP-5MS capillary column (60 m × 0.25 mm i.d., 0.25 μm film thickness). Helium was used as the carrier gas at a constant flow rate of 1.0 mL min^-1^. The inlet was operated in splitless mode. The oven temperature was initially held at 40 °C for 3 min, increased to 180 °C at 3.5 °C min^⁻¹^, and then increased to 310 °C at 10 °C min^⁻¹^. The MS transfer line, ion source, and quadrupole temperatures were set to 240, 230, and 150 °C, respectively. Mass spectra were acquired using electron ionization at approximately 70 eV in full-scan mode over an m/z range of 29-500, with a solvent delay of 3 min. Compounds were provisionally annotated by comparison of their mass spectra with the NIST mass spectral library (NIST23). Identification based solely on library matching was considered tentative. Each sample type was analyzed using three independent biological replicates.

### 5.3. Identification, curation and sequence analysis of D. nobile TPS genes

Thirty-three putative *TPS-a* subfamily genes were retrieved from the chromosome-level*D. nobile* genome reported previously (Li et al., 2024). Gene structures were manually curated using GSAman (Chen et al., 2026). Sequences encoding complete open reading frames with the expected class I terpene synthase domains were retained. The physicochemical properties of proteins were analyzed using ExPASy ProtParam (Gasteiger et al., 2005). DnTPS protein sequences were aligned using CLC sequence. Phylogenetic trees were constructed using the maximum-likelihood (ML) method based on the aligned protein sequences. Branch support was assessed by bootstrap analysis with 5,000 replicates. The resulting trees were visualized and annotated using EvolView. Conserved motifs were identified with MEME Suite implemented in TBtools (Bailey et al., 2015; Chen et al., 2023).

### 5.4. Cloning, heterologous expression, and GC-MS screening

The full-length coding sequences of the curated *DnTPS* genes were amplified from pooled cDNA prepared from different tissues of *D. nobile* using I5 high-fidelity DNA polymerase (Tsingke Biotechnology, China) and gene-specific primers (Supplementary Table S3). The amplified fragments were inserted into the pET28a expression vector at the EcoRI and SalI restriction sites. Each resulting construct was co-transformed into *Escherichia coli* C41 with the FPP-supplying plasmid pMEVT-MBIS. Transformants were cultured in Luria-Bertani (LB) medium supplemented with 50 μg/mL kanamycin and 50 μg/mL chloramphenicol at 37 °C with shaking at 180 rpm. When the cultures reached an OD_600_ of 0.6-0.8, isopropyl β-D-1-thiogalactopyranoside (IPTG) and MgCl_2_ were added to final concentrations of 1 mM and 1 mM MgCl_2_, respectively. Cultures were subsequently incubated at 16 °C with shaking for 3 days.

Following induction, an equal volume of n-hexane was added to each culture. The mixtures were sonicated for 30 min and then left to stand overnight. The upper organic phase was collected and evaporated to dryness using a rotary evaporator. The residues were redissolved in 1 mL of chromatography-grade n-hexane and transferred to ampoules for subsequent GC-MS analysis. Cells harboring the empty pET28a vector together with pMEVT-MBIS served as negative controls. GC-MS analysis was performed using a Shimadzu gas chromatography-mass spectrometry system equipped with a SH-Rxi-5Sil MS column (30 m×0.25 mm i.d., 0.25 μm film thickness). The inlet was operated in splitless mode. The oven temperature was initially held at 100 °C for 3 min, increased to 240 °C at 10 °C min^⁻¹^, and then increased to 300 °C at 20 °C min^⁻¹^, at 300 °C for 3 min, giving a total run time of 23 min. The remaining GC-MS conditions were the same as those described in Section 5.2.

### 5.5. Preparative isolation and NMR characterization of copaborneol

The strain co-expressing *DnTPS1* and the FPP-supplying pathway was cultured at a preparative scale of 6 L. Products were extracted with n-hexane and concentrated. The crude extract was fractionated by normal-phase silica gel column chromatography using an n-hexane/ethyl acetate gradient of 100:1, 80:1, 60:1, 40:1, 20:1, and 10:1 (v/v). Fractions were analyzed by GC-MS to locate the target compound. Target-containing fractions were further purified by silica gel column chromatography using an n-hexane/acetone gradient of 10:1, 9:1, 8:1 and 6:1 (v/v). Purified 3 mg copaborneol was dissolved in CDCl₃ and transferred to an NMR tube. One-dimensional NMR spectra were acquired at 22 °C using a 400 MHz Bruker AVANCE spectrometer. Data were processed using TopSpin 2.0 and MestReNova 15.6. Chemical shifts were referenced to CDCl₃. The structure was assigned by interpretation of the NMR spectra and comparison with published spectroscopic data for copaborneol (Lusi et al., 2022).

### 5.6. Recombinant-protein purification and in vitro enzyme assay

Recombinant DnTPS1 was expressed in *E. coli BL21* carrying pET28a/*DnTPS1*. Cells were induced under 16 °C for 20 h, collected, resuspended in lysis buffer, and disrupted by ultrasonic cell disruptor. The soluble fraction was purified by Ni-NTA, and protein purity was assessed by SDS-PAGE. Enzyme assays were performed in a total volume of 300 μL containing 10 μg of purified DnTPS1, 23 μM farnesyl diphosphate (FPP; Sigma), and assay buffer (50 mM Tris-HCl, 10 mM MgCl₂, 100 mM KCl, and 5% (v/v) glycerol, pH 7.4.). Reaction mixtures were gently mixed and incubated in a water bath at 30 °C, 12 h. Products were extracted with 1 mL of n-hexane by vortexing for 1 min. The organic phase was collected and evaporated to dryness under a stream of nitrogen. The residue was redissolved in 100 μL of n-hexane and transferred to a GC vial for analysis using GC-MS conditions the same as those described in Section 5.4. The product was compared with purified copaborneol.

### 5.7. Transient expression and subcellular localization

For transient expression in *Nicotiana benthamiana*, the *DnTPS1* coding sequence was inserted into pCAMBIA1300 by homologous recombination. The resulting pCAMBIA1300/*DnTPS1* construct together with the *tHMGR* was introduced into *Agrobacterium tumefaciens EHA105* competent cells using the freeze-thaw method. The bacterial suspension was adjusted to an OD₆₀₀ of 0.4-0.6 using infiltration buffer containing 100 μM acetosyringone and 10 mM MgCl₂. The suspension was pressure-infiltrated into tobacco leaves using a needleless syringe. Plants were maintained for 4-5 days after infiltration. The fresh samples were used for HS-SPME-GC-MS analysis. The GC-MS analysis conditions were the same as those described in Section 5.2.

For subcellular localization, the DnTPS1 coding sequence without the stop codon was fused to eGFP in pAN580. The fusion construct and free-eGFP control were introduced into protoplasts prepared from etiolated rice seedlings by PEG-mediated transfection and transiently expressed in *N. benthamiana* leaves via Agrobacterium-mediated infiltration. Following infiltration, fluorescence signals were observed using an inverted fluorescence microscope. Images were acquired with appropriate filter sets for the corresponding fluorescent proteins, and subcellular localization patterns were determined based on the distribution of fluorescence signals. Localization predictions were obtained using TargetP and WoLF PSORT.

### 5.8. Structural modeling, ligand docking and site-directed mutagenesis

The protein structure of DnTPS1 was predicted using AlphaFold3. The receptor protein was prepared using PyMOL 2.5.2 by removing water molecules, salt ions, and other small molecules. The 3D structure of 2E,6E-FPP was obtained from the PubChem database and subjected to energy minimization under the MMFF94 force field. Molecular docking was performed using AutoDock Vina 1.2.3, with the docking box set to encompass the entire protein. The prepared ligand and receptor were converted into PDBQT format using ADFRsuite 1.0. Finally, docking poses were visualized and analyzed using PyMOL 2.5.2 to identify key amino acid residues located near the predicted active pocket.

Multiple sequence alignment was performed using CLC Sequence Viewer 8.0. The sequences analyzed included DnTPS1, DnTPS2, and DnTPS3, previously reported plant copaborneol-producing sesquiterpene synthases, including *ZmTPS8* (Saldivar et al., 2023) and *MtTPS5* (Garms et al., 2010), and other functionally characterized sesquiterpene synthases. Based on the alignment, residues located within the catalytic cavity of DnTPS1 and showing marked differences in amino acid properties compared with the corresponding positions in other enzymes were selected, including Y12, I295, C298, W377, T402, T403, and C404. Using pET28a/*DnTPS1* as the template, site-directed mutagenesis was performed according to the instructions of the Stratagene QuikChange Kit to generate the Y12F, I295A, I295L, C298A, C298L, W377Y, W377F, T402S, T403A, T403C, C404A, and C404G variants. The catalytic products of the mutant enzymes were subsequently analyzed after heterologous expression in *E. coli* by the GC-MS procedure described in Section 5.4.

### 5.9. Comparative phylogenetic and local gene-organization analyses

Using the BLAST function in TBtools, protein sequences highly homologous to DnTPS1 were identified from the genomes of five Dendrobium species. Sequences shorter than 300 amino acids were excluded, and a total of 15 TPS sequences were retained for subsequent analysis. TPS proteins were aligned with MAFFT v7.526 (Katoh and Standley, 2013) using the L-INS-i strategy (--localpair --maxiterate 1000) and trimmed with ClipKIT v2.13.2 (Steenwyk et al., 2020) in smart-gap mode, retaining 410 amino-acid sites. A maximum-likelihood tree was inferred in IQ-TREE v2.4.0 (Minh et al., 2020) under Q.plant+G4 selected by ModelFinder using BIC. Branch support was estimated with 1,000 SH-aLRT replicates and 1,000 ultrafast bootstrap replicates with -bnni.

Pairwise local alignments of protein sequences within the selected genomic regions were performed using the Smith-Waterman algorithm with the BLOSUM62 substitution matrix. Gap-opening and gap-extension penalties were set to −11 and −1, respectively. Matches were retained if they had an alignment score ≥100, at least 80 aligned positions excluding gaps, sequence identity ≥35%, and bidirectional sequence coverage ≥50%. Reciprocal best hits (RBHs) within each genomic region pair were then identified based on alignment scores. According to the chromosomal order of genes, ordered chains of RBH anchors were constructed between adjacent displayed genomic regions. Each chain was required to contain at least three anchor pairs, with no more than 10 annotated genes between consecutive anchors. For each pair of genomic regions, the longest anchor chain was selected for visualization. Homologous relationships among TPS candidate genes were identified separately using the same alignment thresholds, while allowing one-to-many matches. The partial match between Dthyrsi01G00342.1 and Dof09G002520.1, which did not meet the bidirectional coverage threshold, was specifically annotated. Gene positions and orientations were determined based on the GFF annotations, and each genomic region was independently scaled according to its coordinate range.

## CRediT authorship contribution statement

Dandan Li: Conceptualization, Methodology, Supervision, Formal analysis, Data curation, Writing-original draft, Writing-review & editing. Jianling Wang: Investigation (gene cloning, vector construction, and heterologous expression in *E. coli* and *N. benthamiana*). Sijing Fu: Investigation (heterologous expression in *E. coli*). Hualei Wang: Funding acquisition.

## Data availability

The corrected coding sequences of all 16 *DnTPS* genes have been submitted to NCBI GenBank and are awaiting accession number assignment.

## Declaration of generative AI and AI-assisted technologies in the manuscript preparation process

During the preparation of this manuscript, the authors used AI for language editing, translation assistance, and reference formatting. After using this tool, the authors reviewed and edited the content as needed and take full responsibility for the content of the published article.

## Acknowledgements

We gratefully acknowledge Professor Qiang Wang from Sichuan Agricultural University for kindly providing the FPP precursor-producing plasmid pMEVT-MBIS. We are grateful to Professor Meirong Jia from the Institute of Materia Medica, Chinese Academy of Medical Sciences, for guidance on the isolation and purification of compounds.

## Supplementary material

**Fig. S1. Electron-ionization mass spectra associated with GC-MS peaks #1∼#27 in** Fig. 1. The panels are labeled, respectively, as α-cubebene (#1), cyclosativene (#2), α-copaene (#3), diepi-α-cedrene (#4), daucene (#5), β-patchoulene (#6), α-elemene (#7), trans-α-bergamotene (#8), β-caryophyllene (#9), aromadendrene (#10), α-guaiene (#11), patchoulene (#12), α-amorphene (#13), eremophilene (#14), β-sesquisabinene (#15), valencene (#16), α-muurolene (#17), δ-cadinene (#18), farnesene (#19), β-bisabolene (#20), zingiberene (#21), β-sesquiphellandrene (#22), nerolidol (#23), germacrene B (#24), cedrol (#25), copaborneol (#26), dendrobine (#27). . The horizontal axis shows m/z, and the vertical axis shows relative ion intensity (%). All names shown here are putative annotations based on the available mass-spectral evidence.

**Fig. S2. Electron-ionization mass spectra associated with GC-MS peaks #1∼#9 in** Fig. 3. The panels are labeled, respectively, as 10-epi-cubebol (#1), cubebol (#2), copaborneol (#3), daucene (#4), α-gurjunene (#5), α-guaiene (#6), cyclosativene (#7), γ-patchoulene (#8) and α-cadinol (#9). The horizontal axis shows m/z, and the vertical axis shows relative ion intensity (%). Except for the separately NMR-characterized copaborneol (#3), names shown here are putative annotations based on the available mass-spectral evidence.

**Fig. S3. ^1^H NMR spectrum of the purified major product generated by DnTPS1.** The product was isolated from the preparative *E. coli* expression system and dissolved in CDCl₃ for analysis. The horizontal axis shows chemical shift (δ, ppm); the solvent resonance near 7.26 ppm is indicated. The spectrum was interpreted together with the ¹³C NMR spectrum in Fig. S2 and published data to support assignment as copaborneol.

**Fig. S4. ¹³C NMR spectrum of the purified major product generated by DnTPS1. The product was analyzed in CDCl₃.** The horizontal axis shows chemical shift (δ, ppm); the solvent signal near 77.2 ppm is marked. Together with the ¹H NMR spectrum in Fig. S1 and published spectroscopic data, these signals support assignment of the DnTPS1 major product as copaborneol.

**Table S1. Candidate compounds detected in different tissues of Dendrobium nobile**

**Table S2. Physicochemical properties and predicted localization of DnTPS proteins**

**Table S3. Cloning and homology-arm primers for 16 *DnTPS* genes.**

