## Supplementary material for "Functional characterization of terpene synthases from *Dendrobium nobile* provides insights into the biosynthesis of copaborneol, a proposed dendrobine precursor": Fig. S1

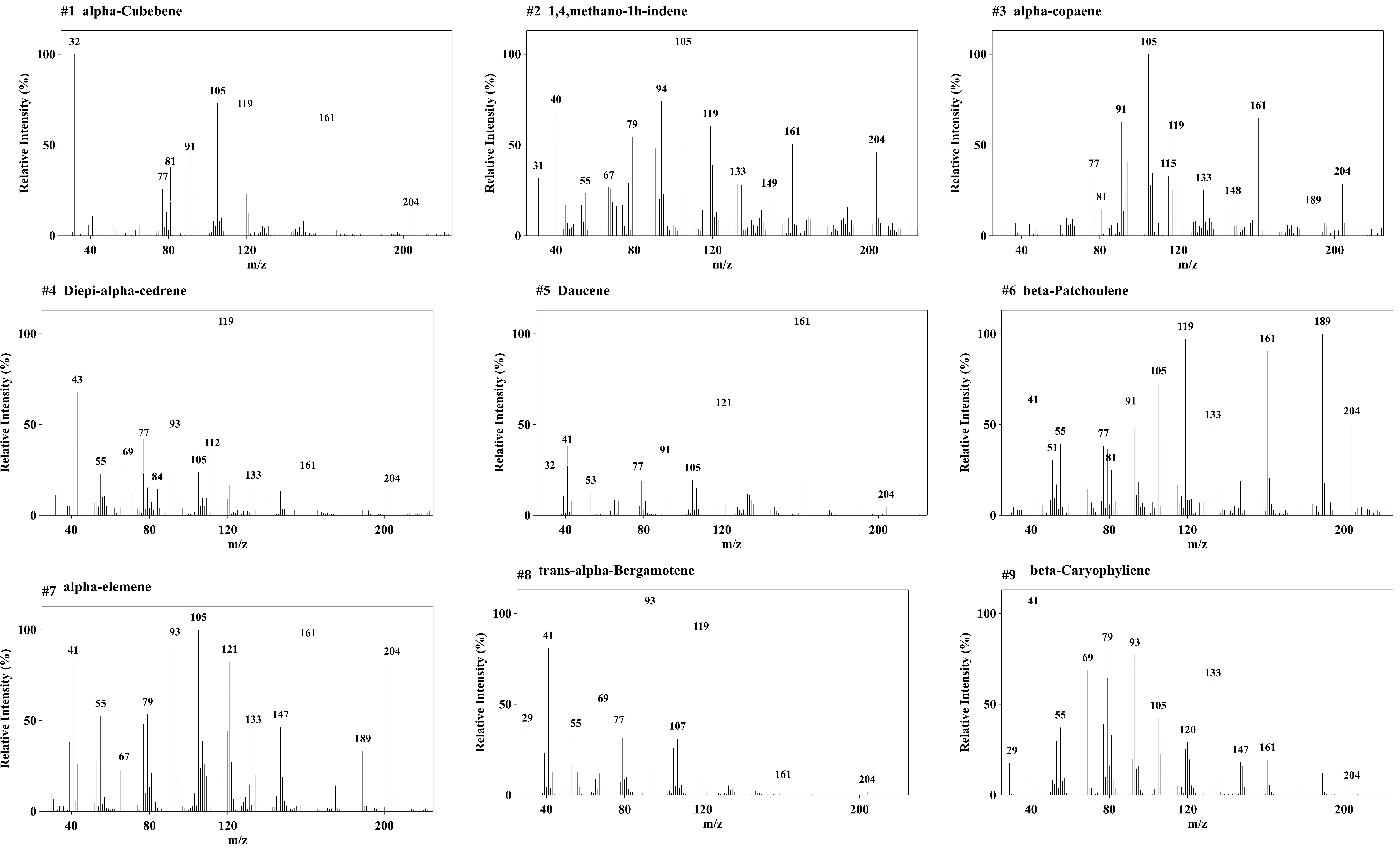
#1 alpha-Cubebene
32
105
119
161
91
81
77
204
#2 1,4,methano-1h-indene
105
40
94
119
79
161
204
31
55
67
133
149
#3 alpha-copaene
105
91
161
119
115
77
81
133
148
189
204
#4 Diepi-alpha-cedrene
119
43
77
93
69
112
55
105
84
133
161
204
#5 Daucene
161
121
41
91
32
53
77
105
204
#6 beta-Patchoulene
119
189
161
105
41
91
55
51
77
81
133
204
#7 alpha-elemene
105
93
121
161
204
41
55
79
67
133
147
189
#8 trans-alpha-Bergamotene
93
119
41
29
69
55
77
107
161
204
#9 beta-Caryophyliene
41
79
93
69
133
105
55
120
29
147
161
204
Relative Intensity (%)
m/z
0
50
100
40
80
120
200

**#10 Aromadendrene**

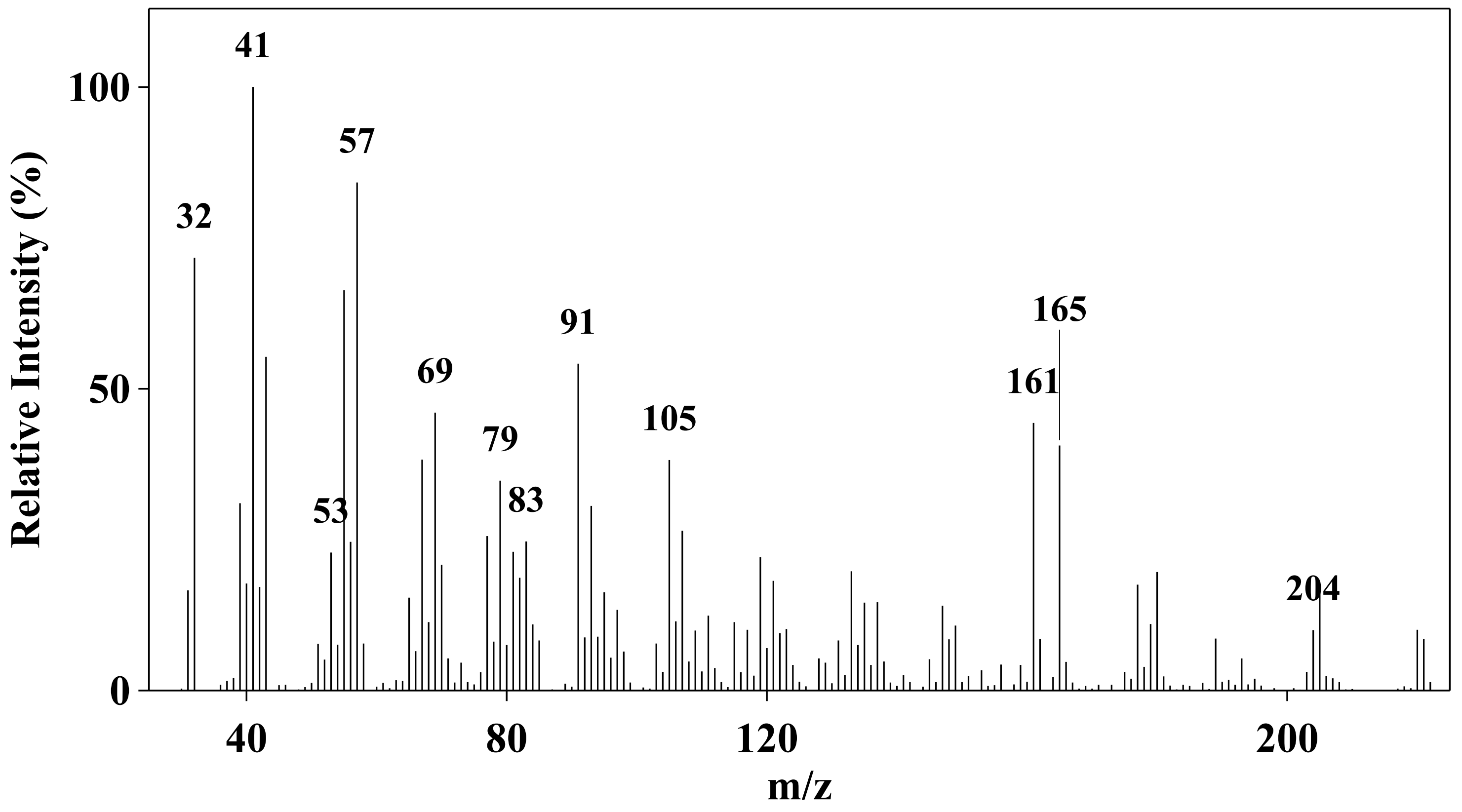


**#11 alpha-Guaiene**

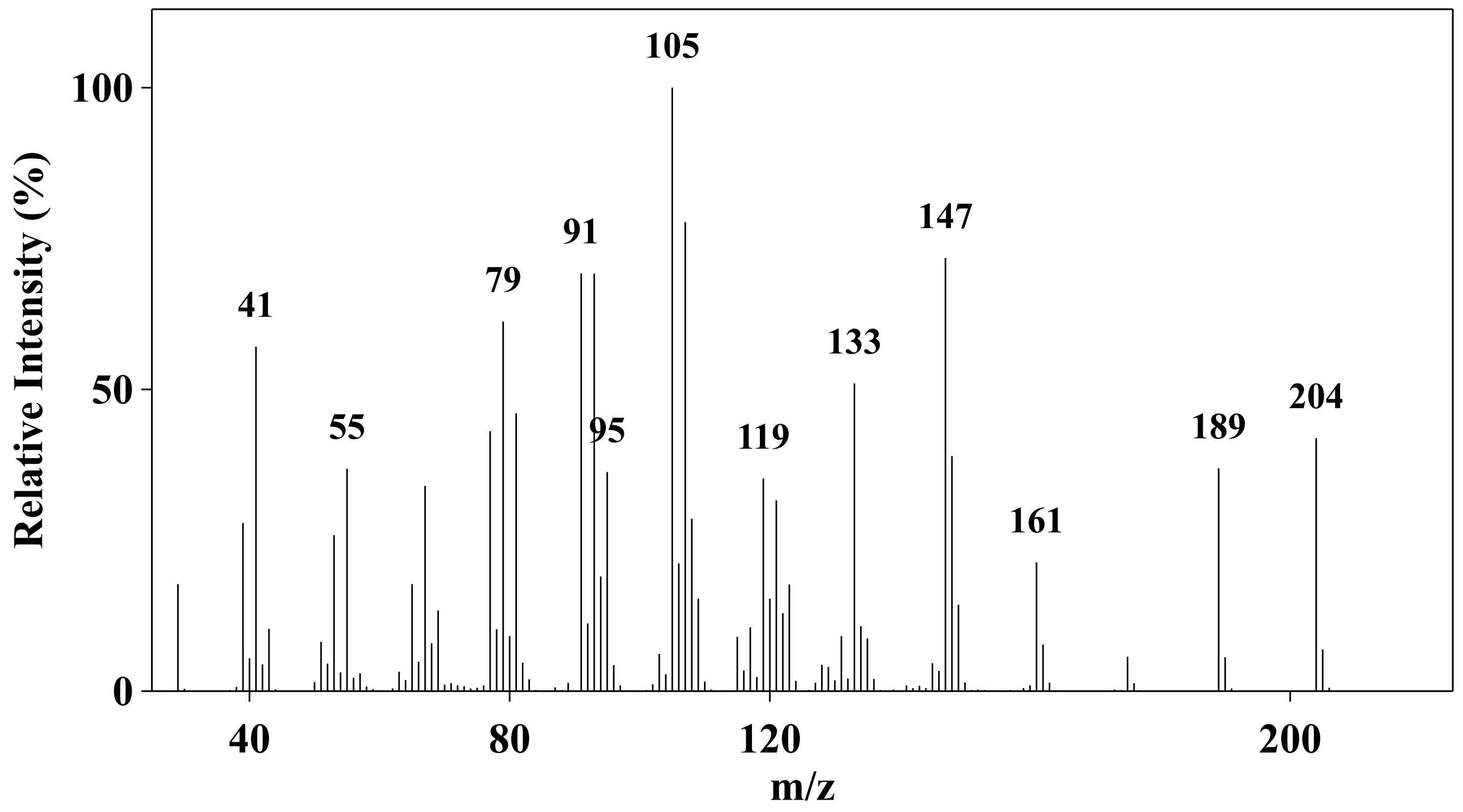


**#12 patchoulene**

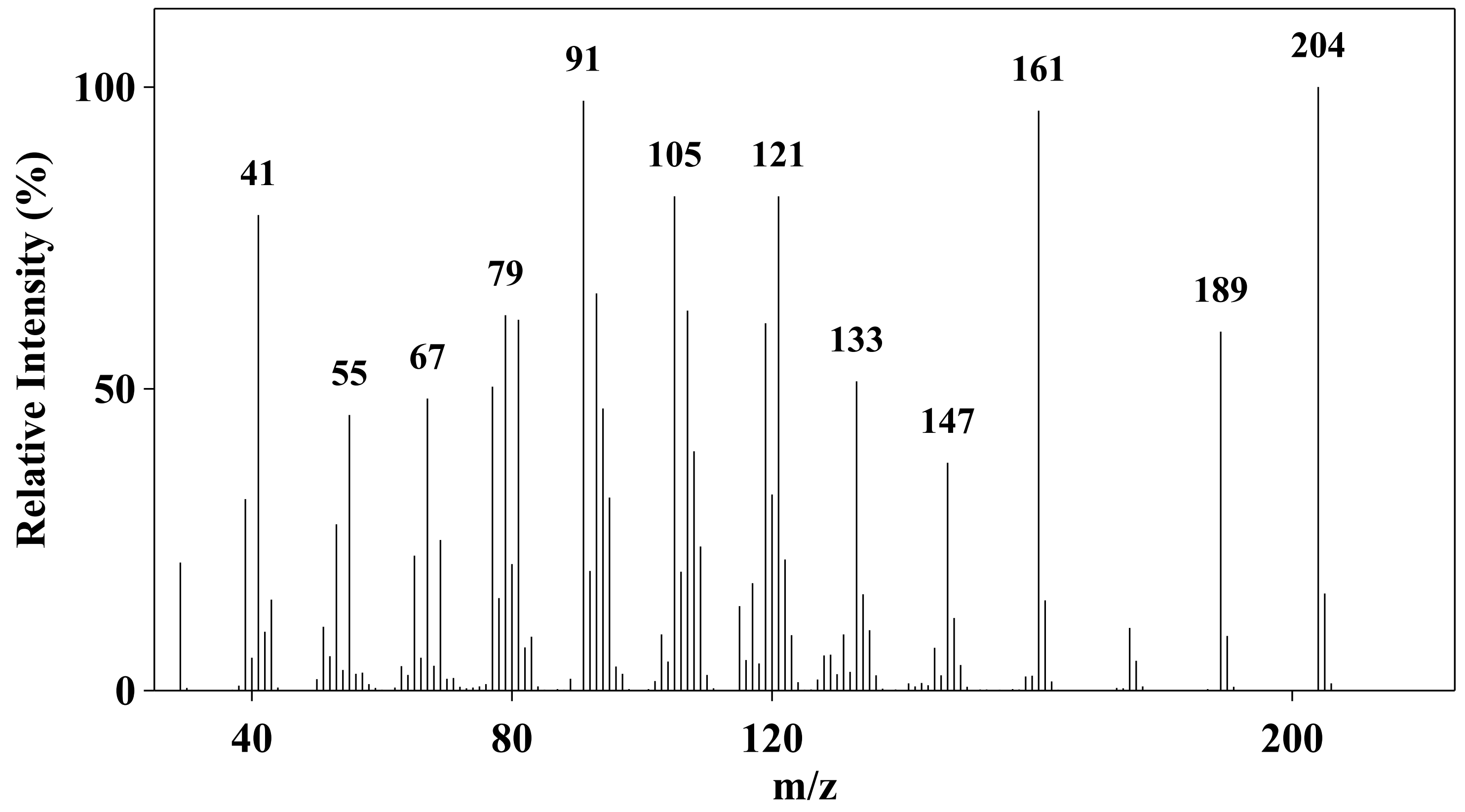


**#13 alpha-amorphene**

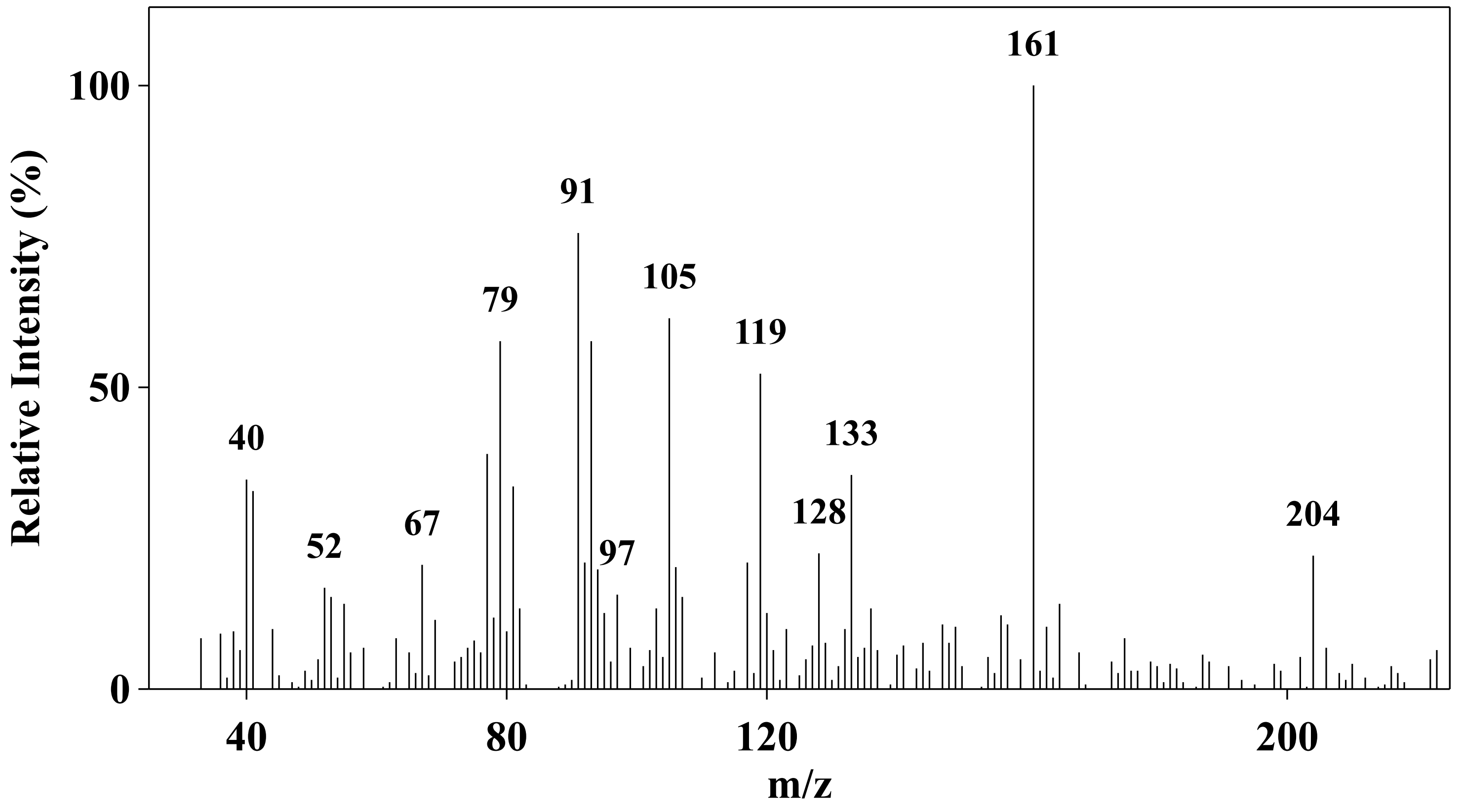


**#14 eremophilene**

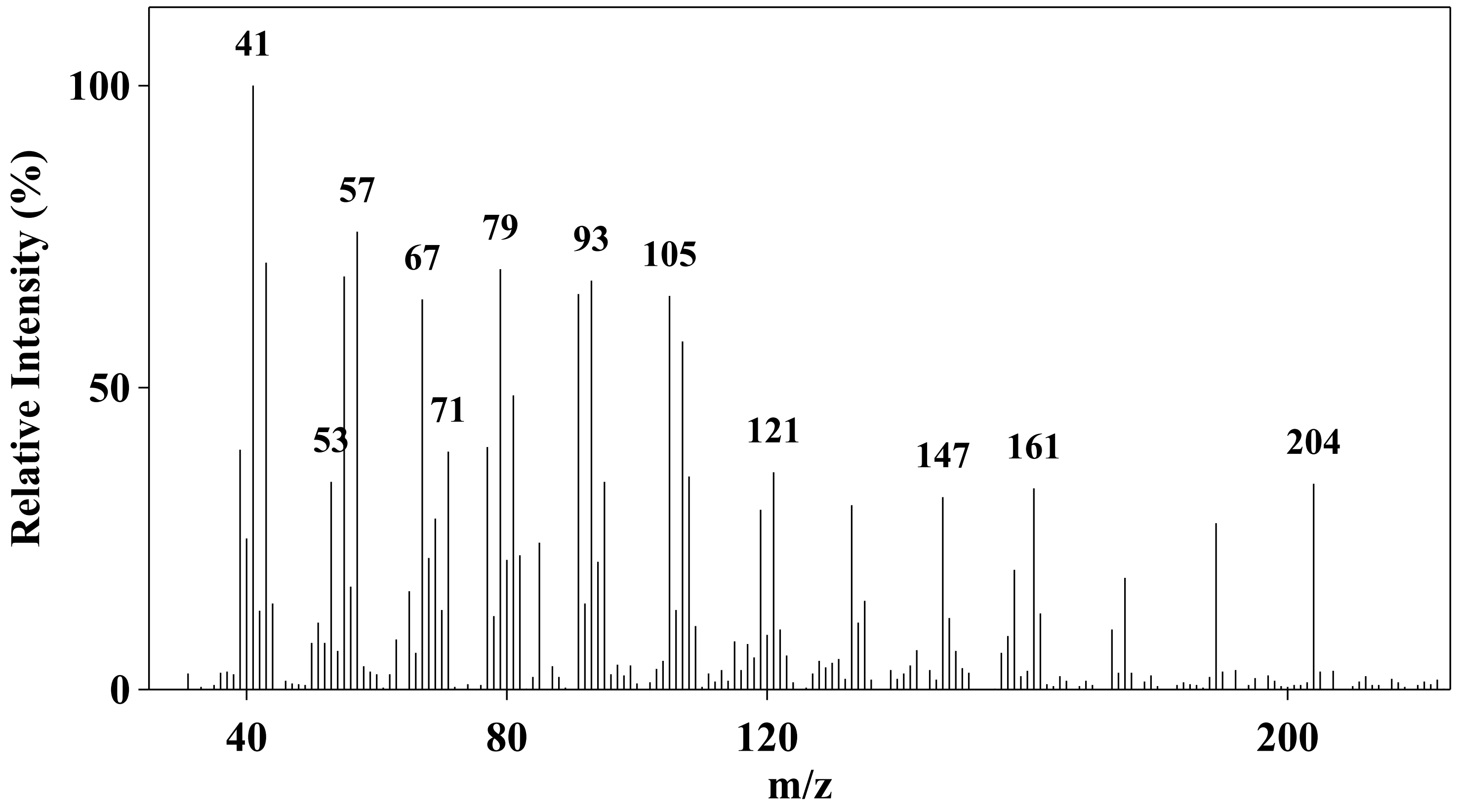


**#15 beta-Sesquisabinene**

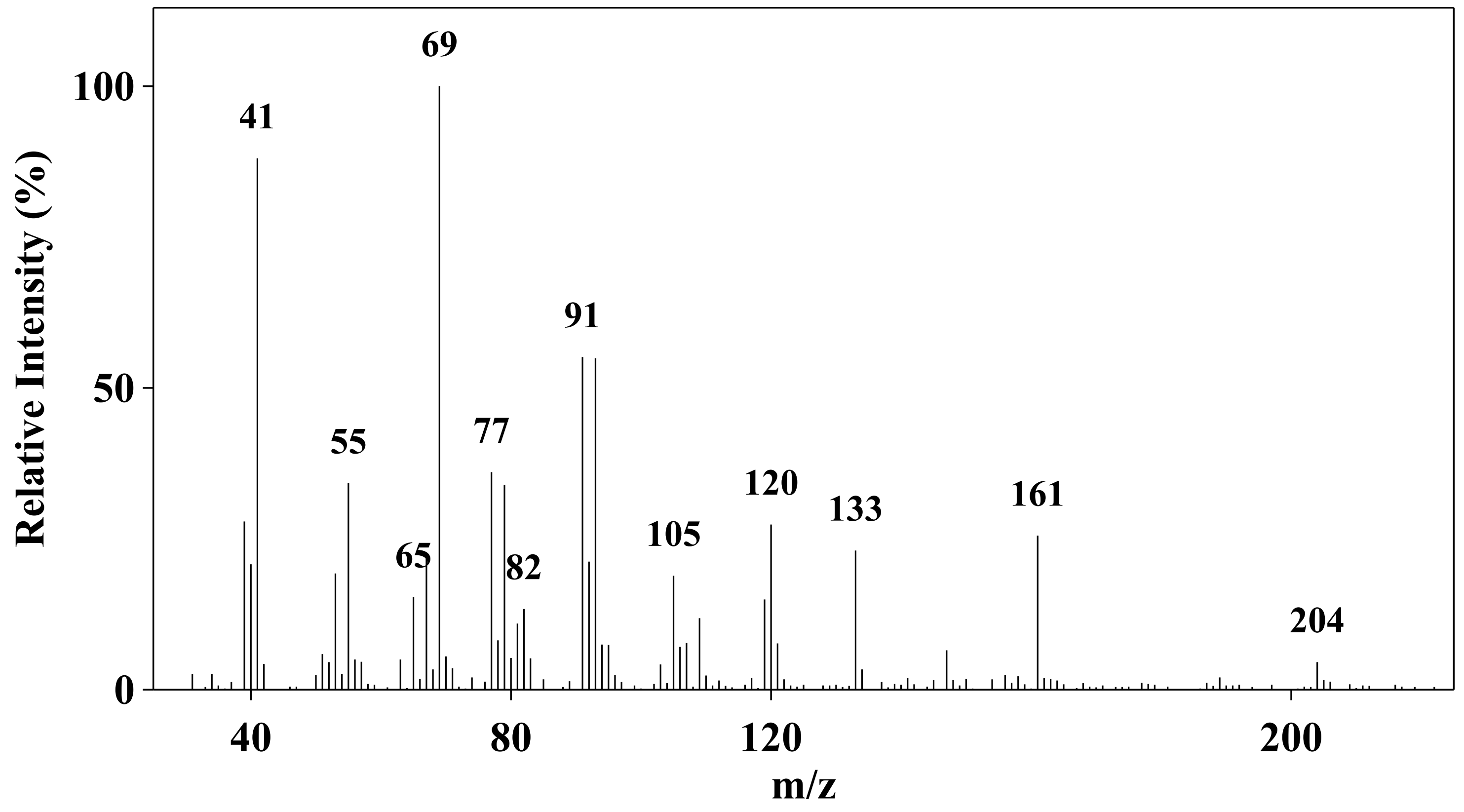


**#16 valencene**

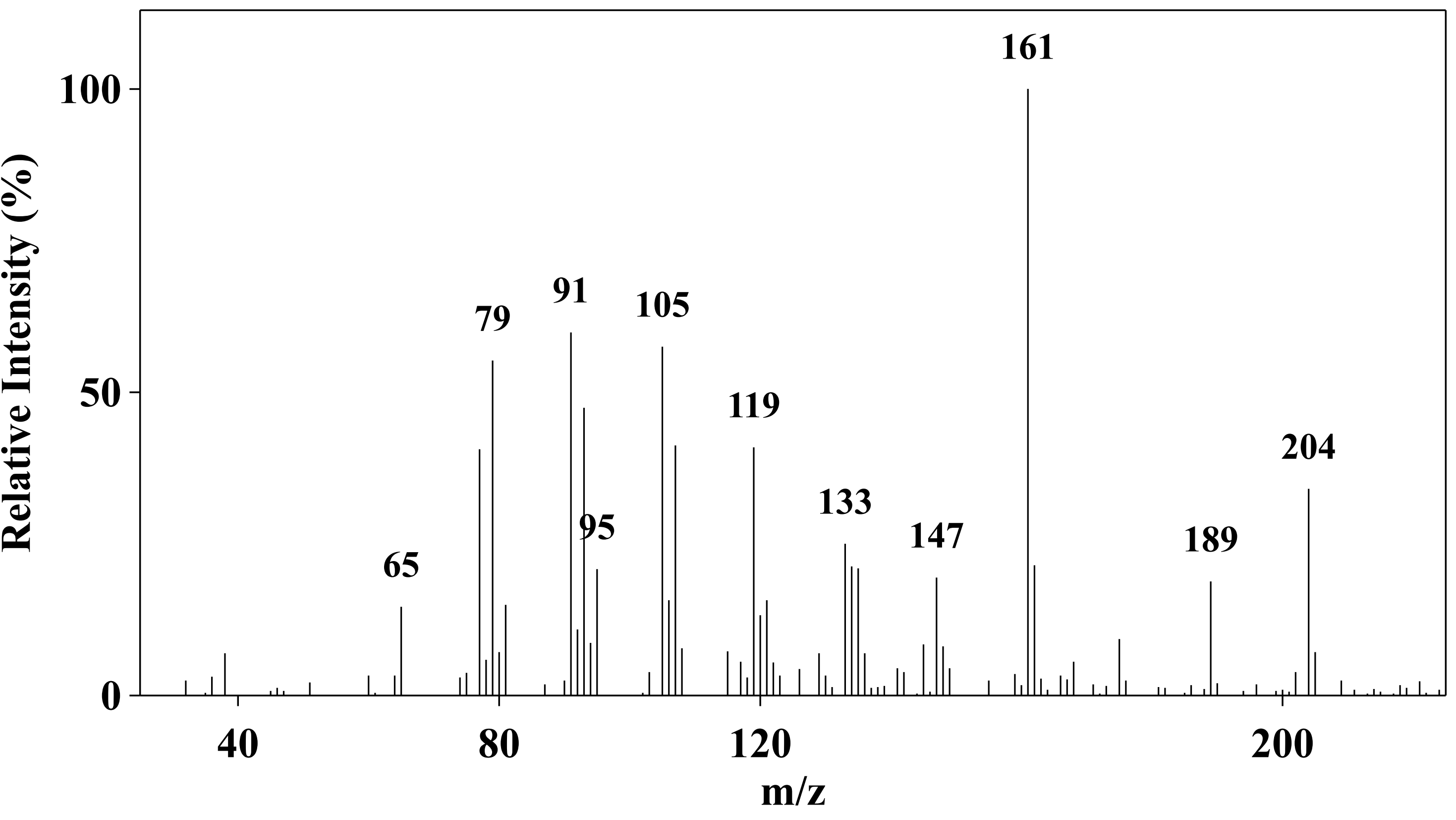


**#17 alpha-Muurolene**

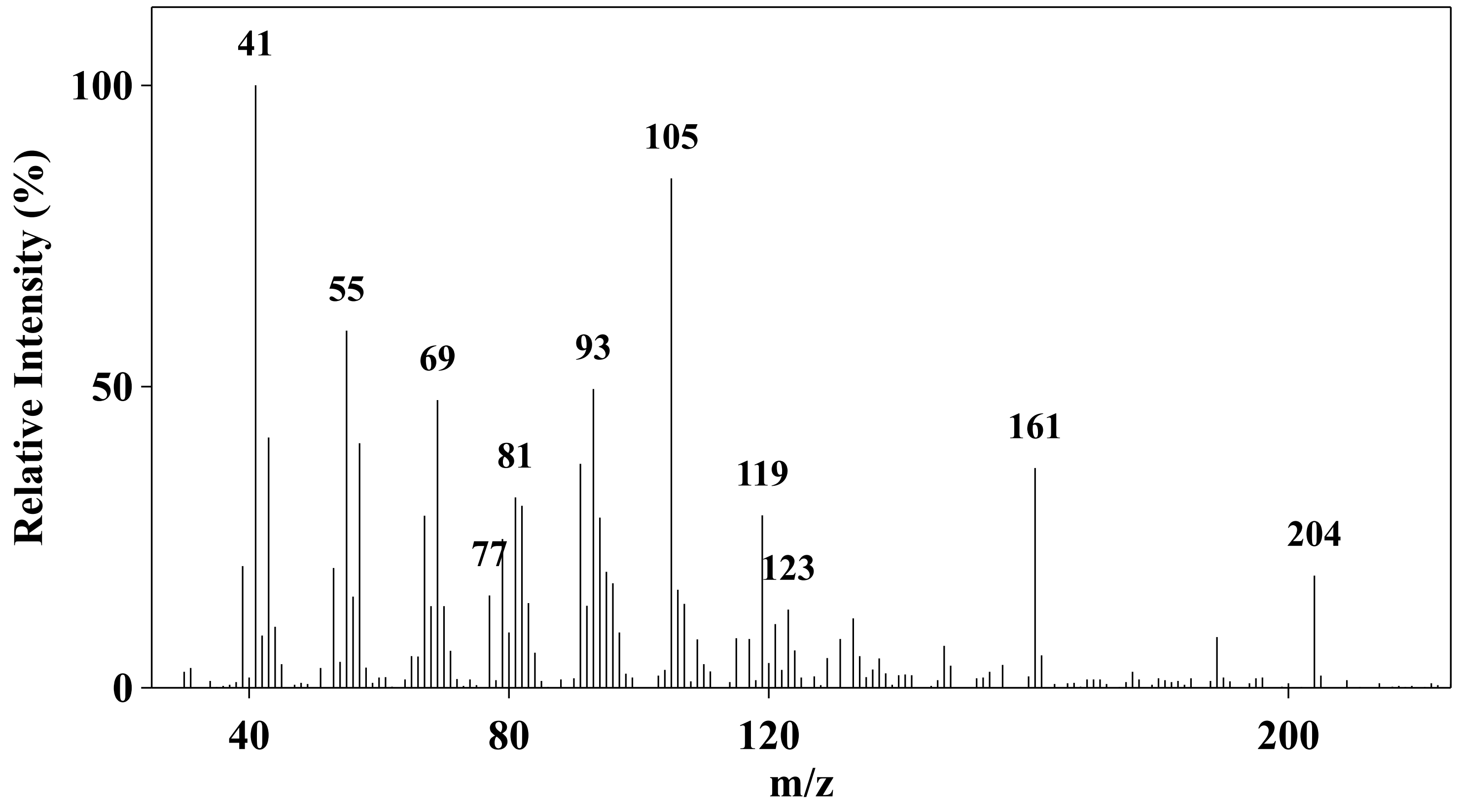


**#18 deta-cadinene**

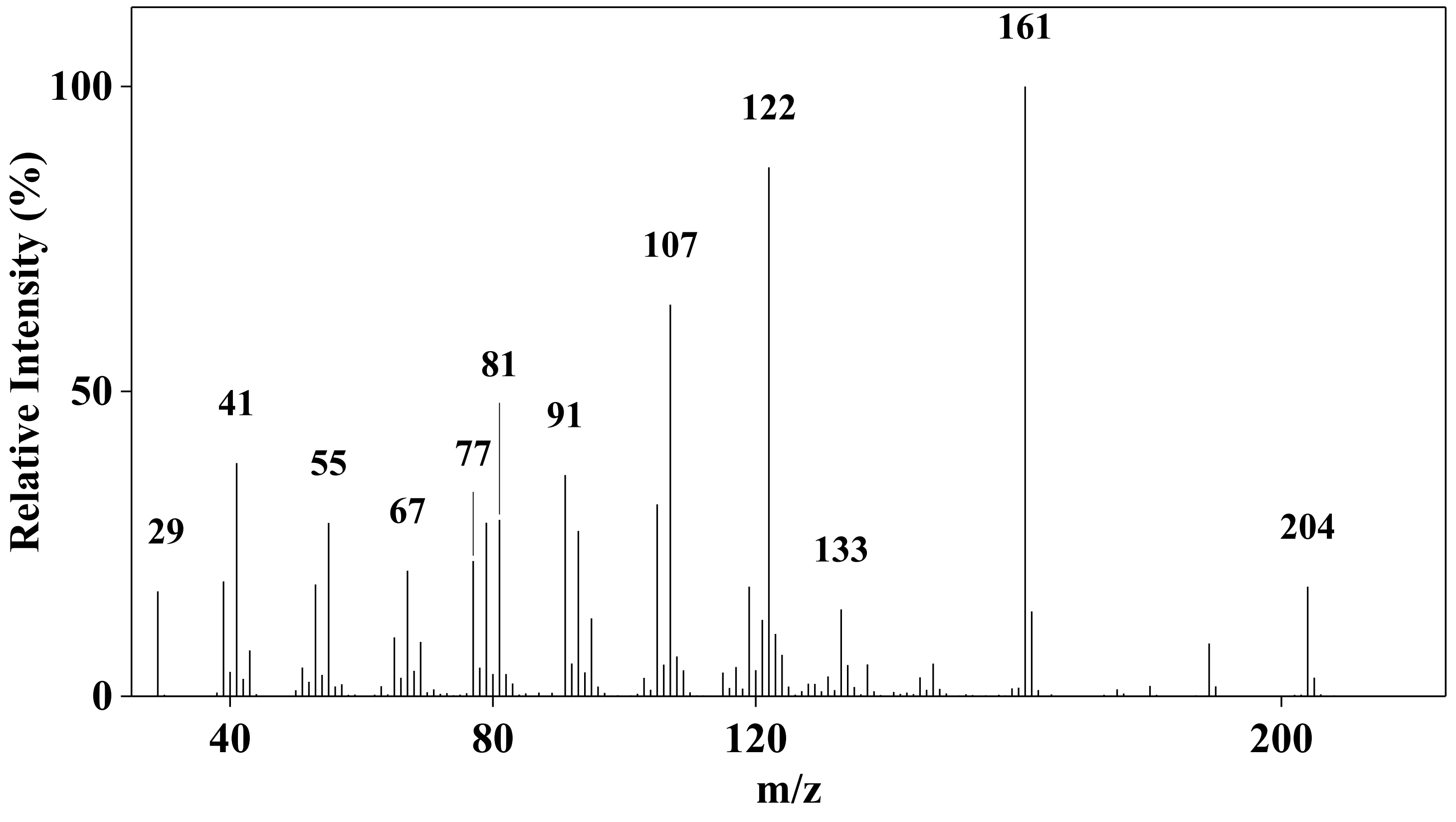

**#19 farnesene**

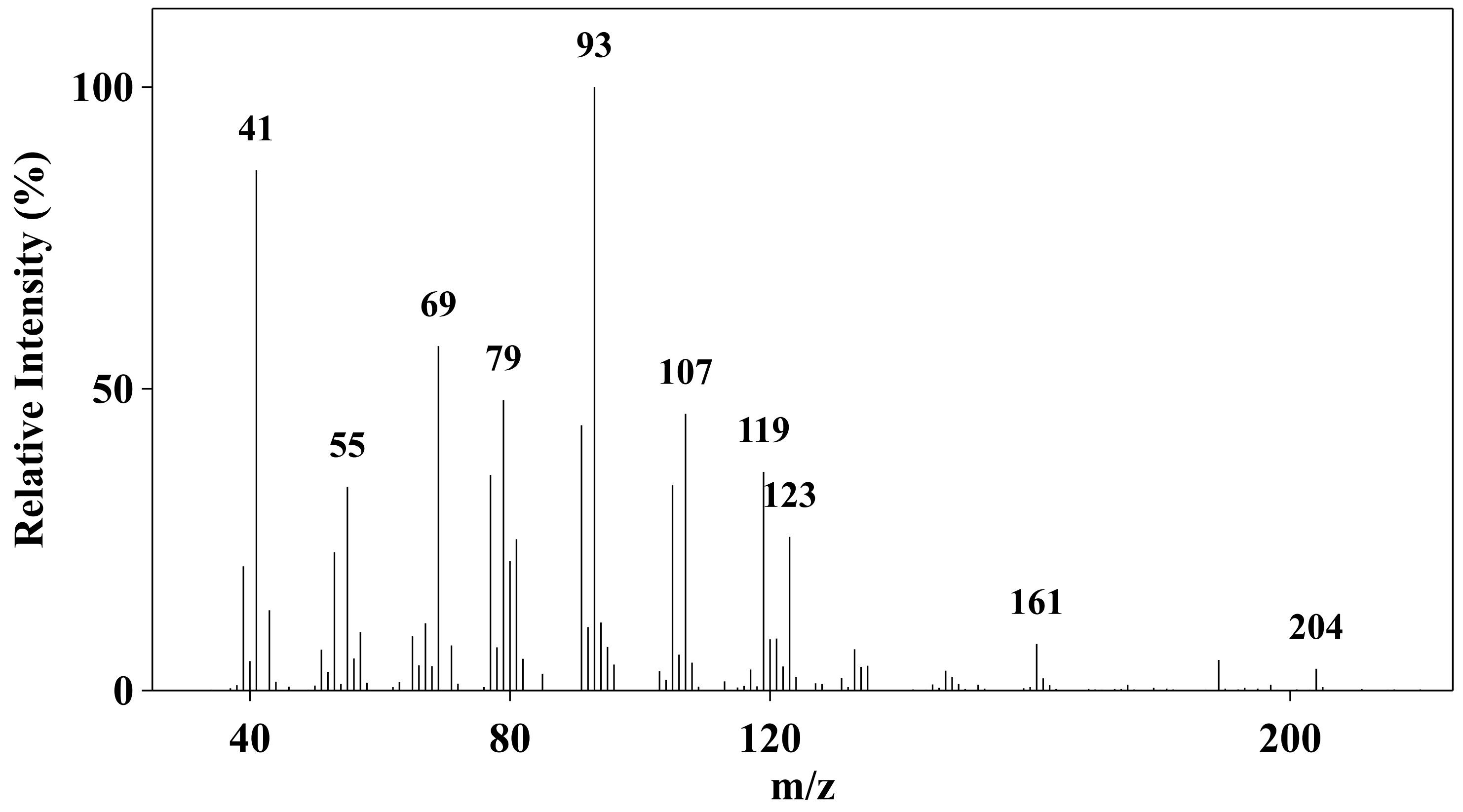


**#20 beta-Bisabolene**

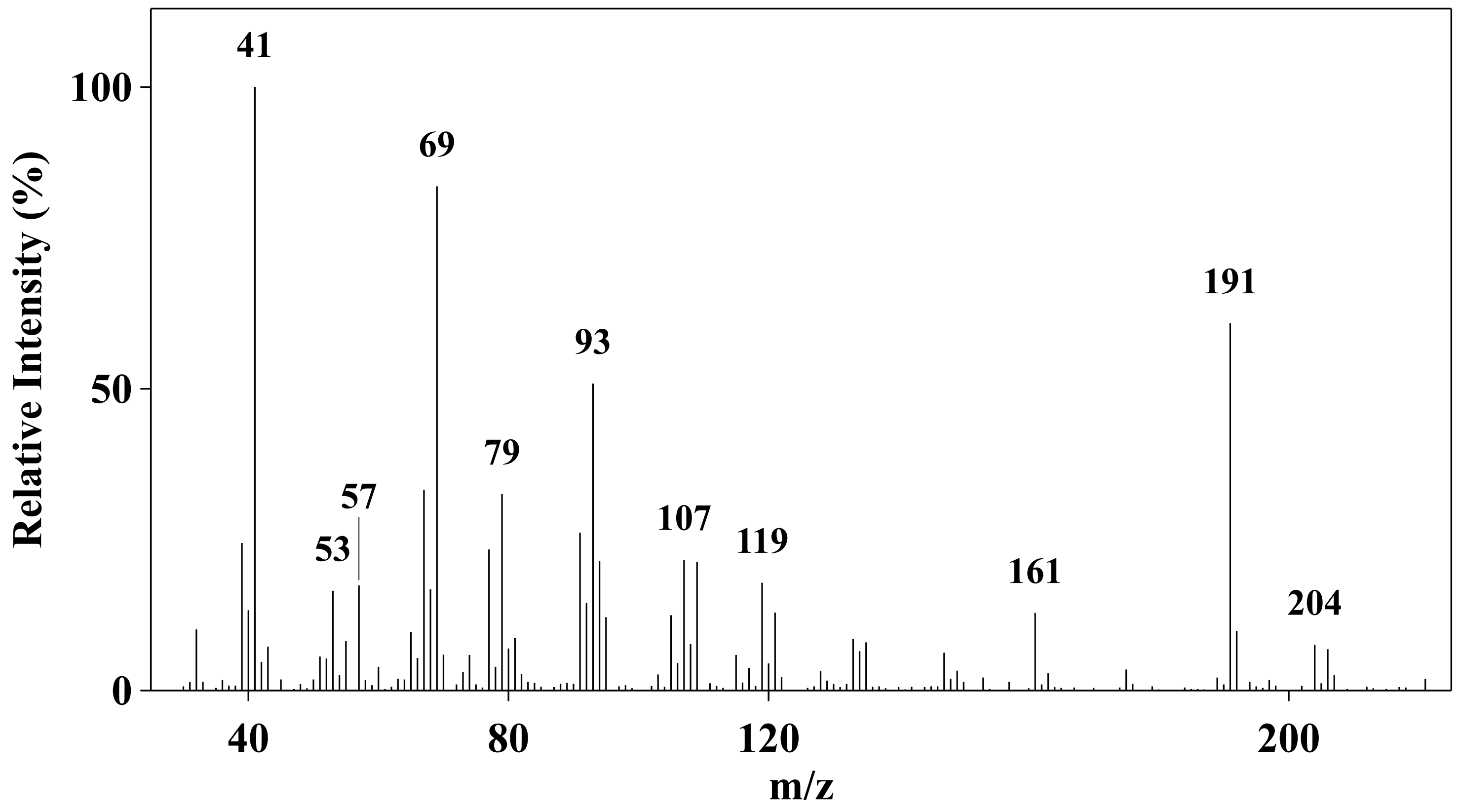


**#21 zingiberene**

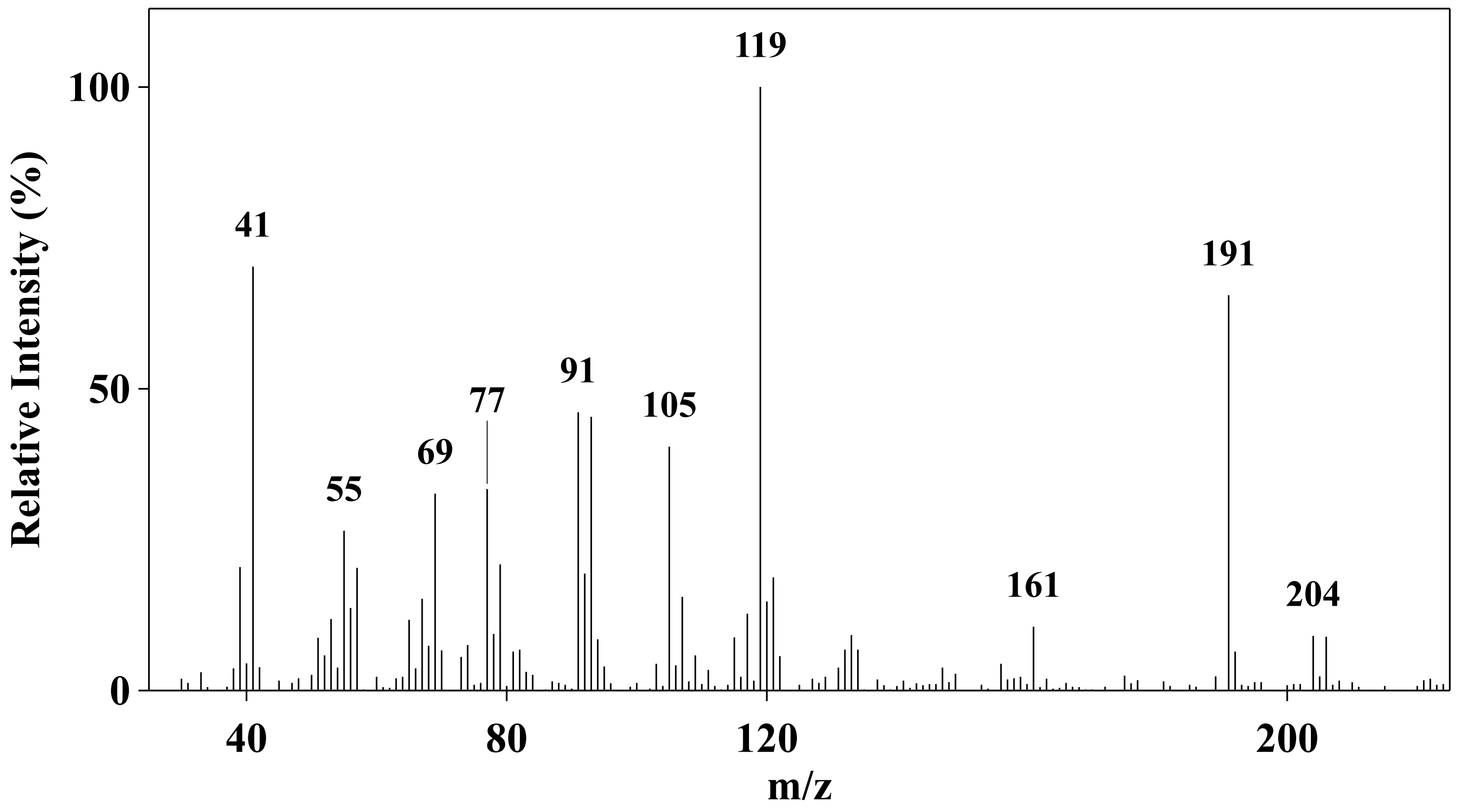


**#22 beta-sequiphellandrene**

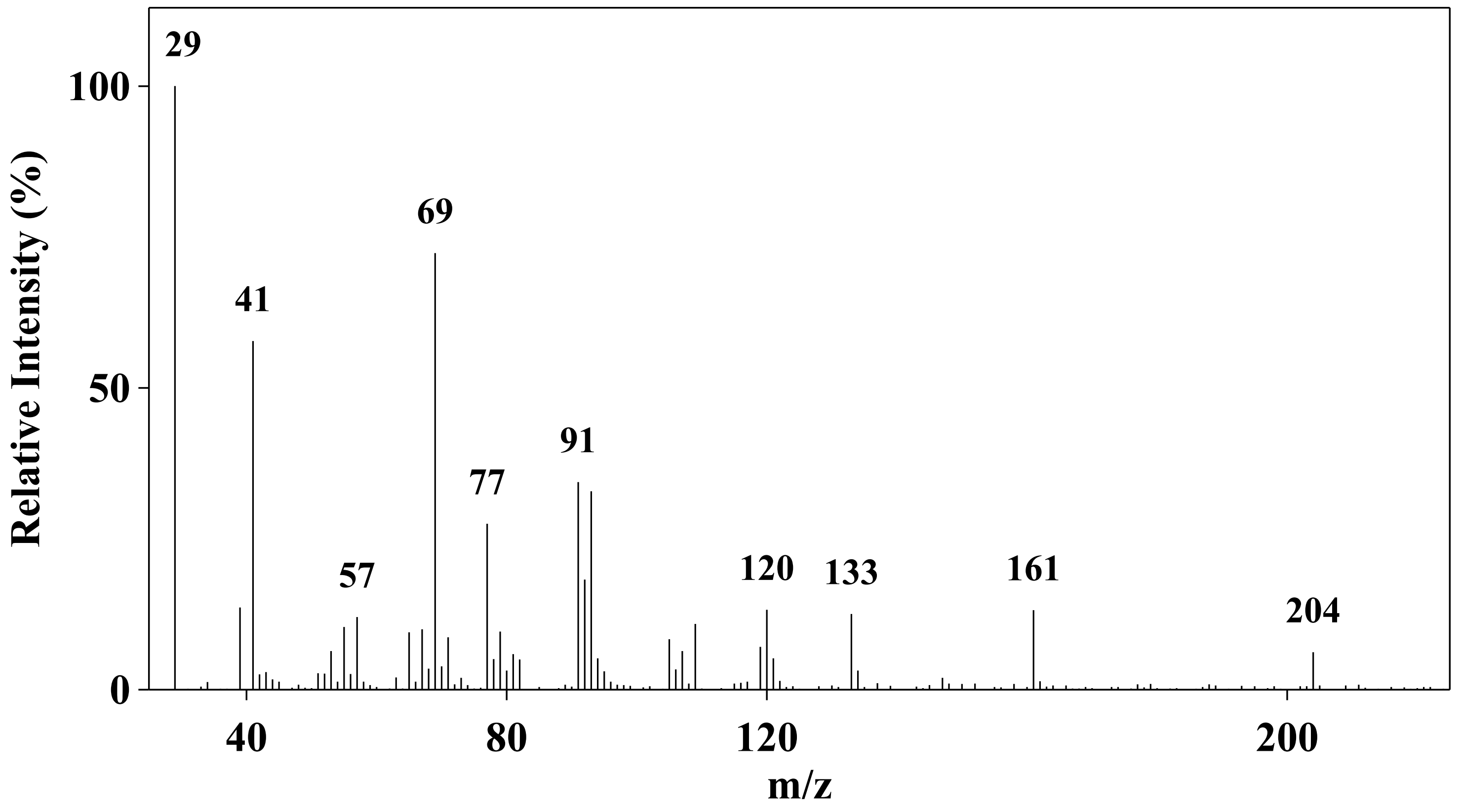


**#23 1,6,10-dodecatrien-ol**

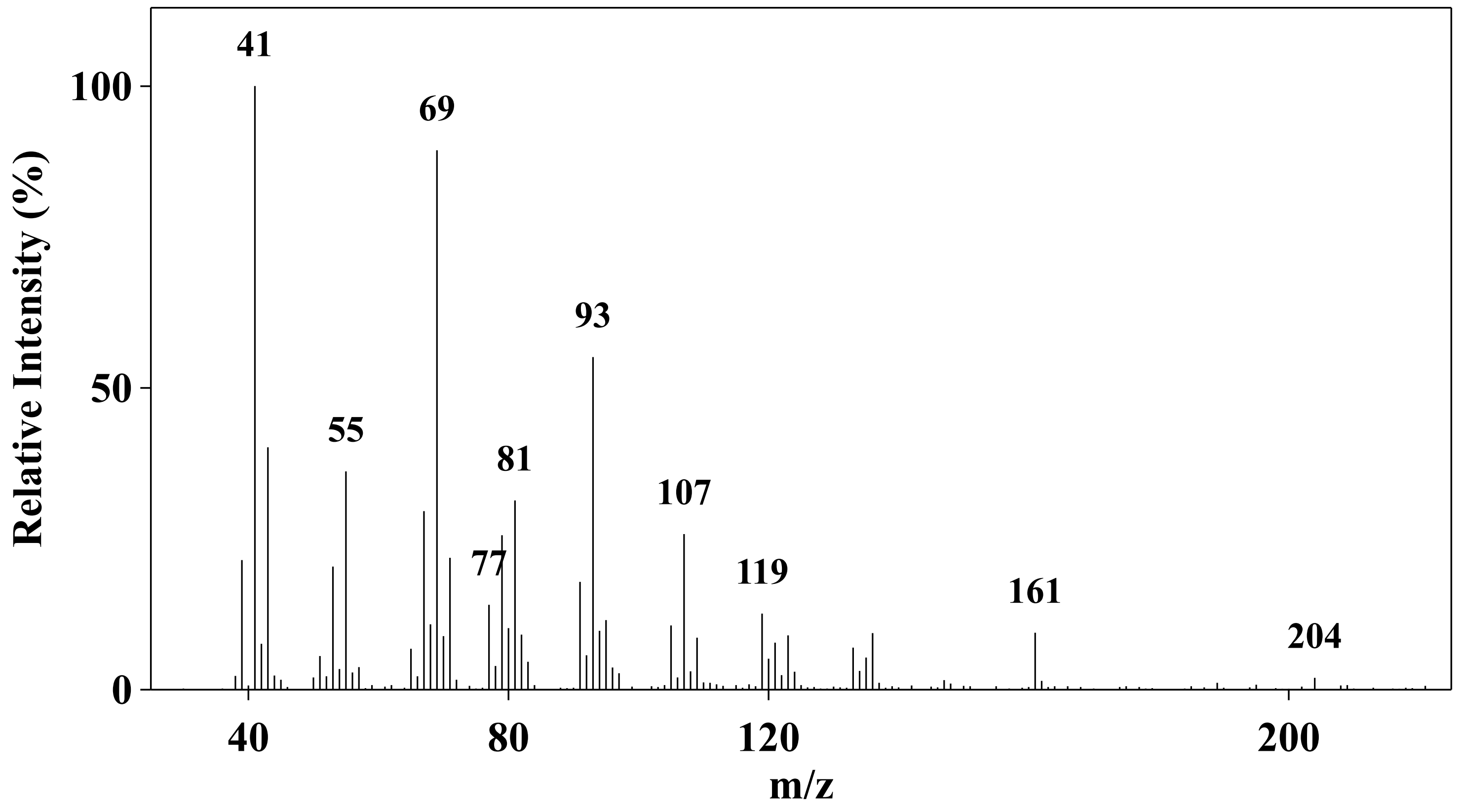


**#24 Germacrene B**

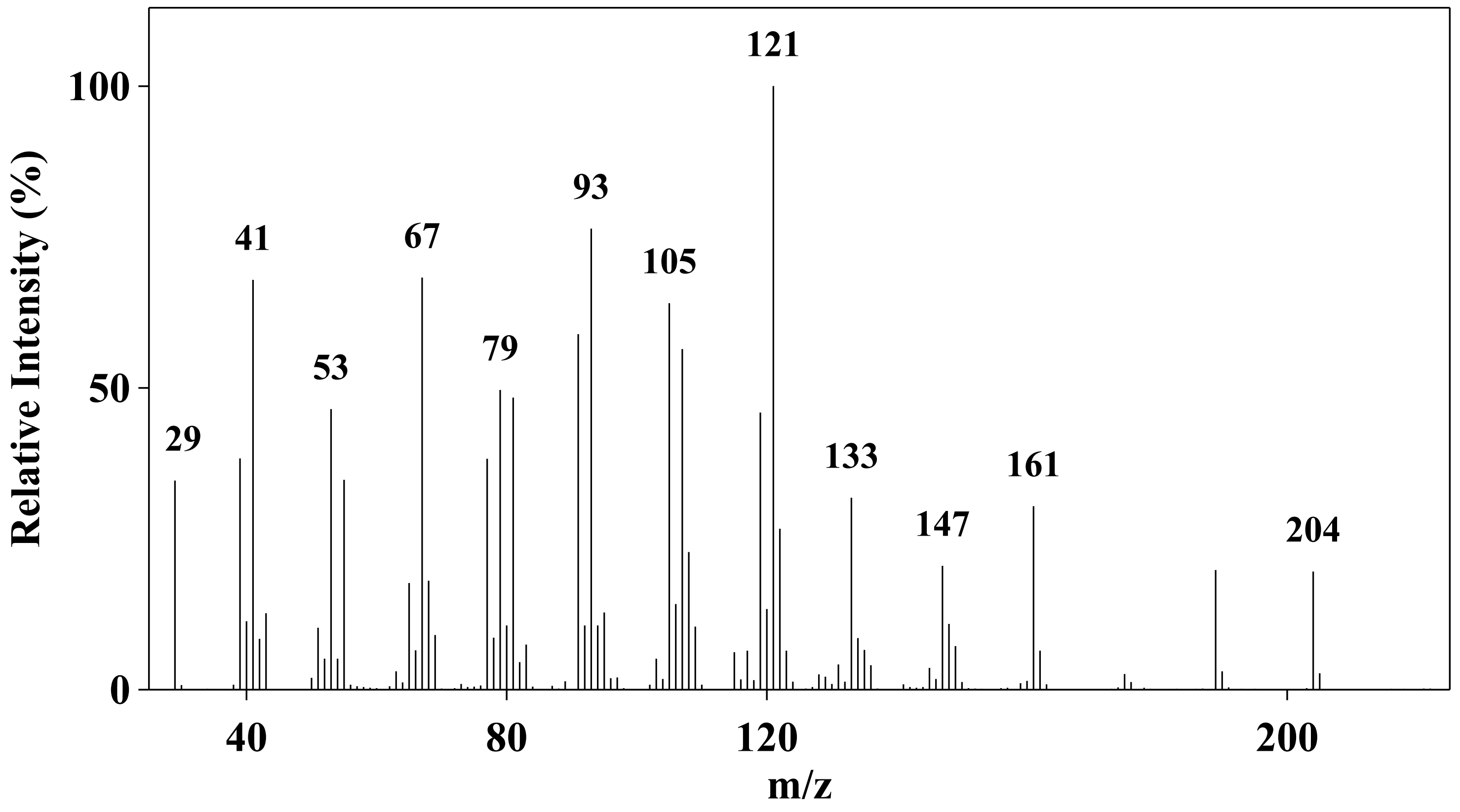


**#25 Cedrol**

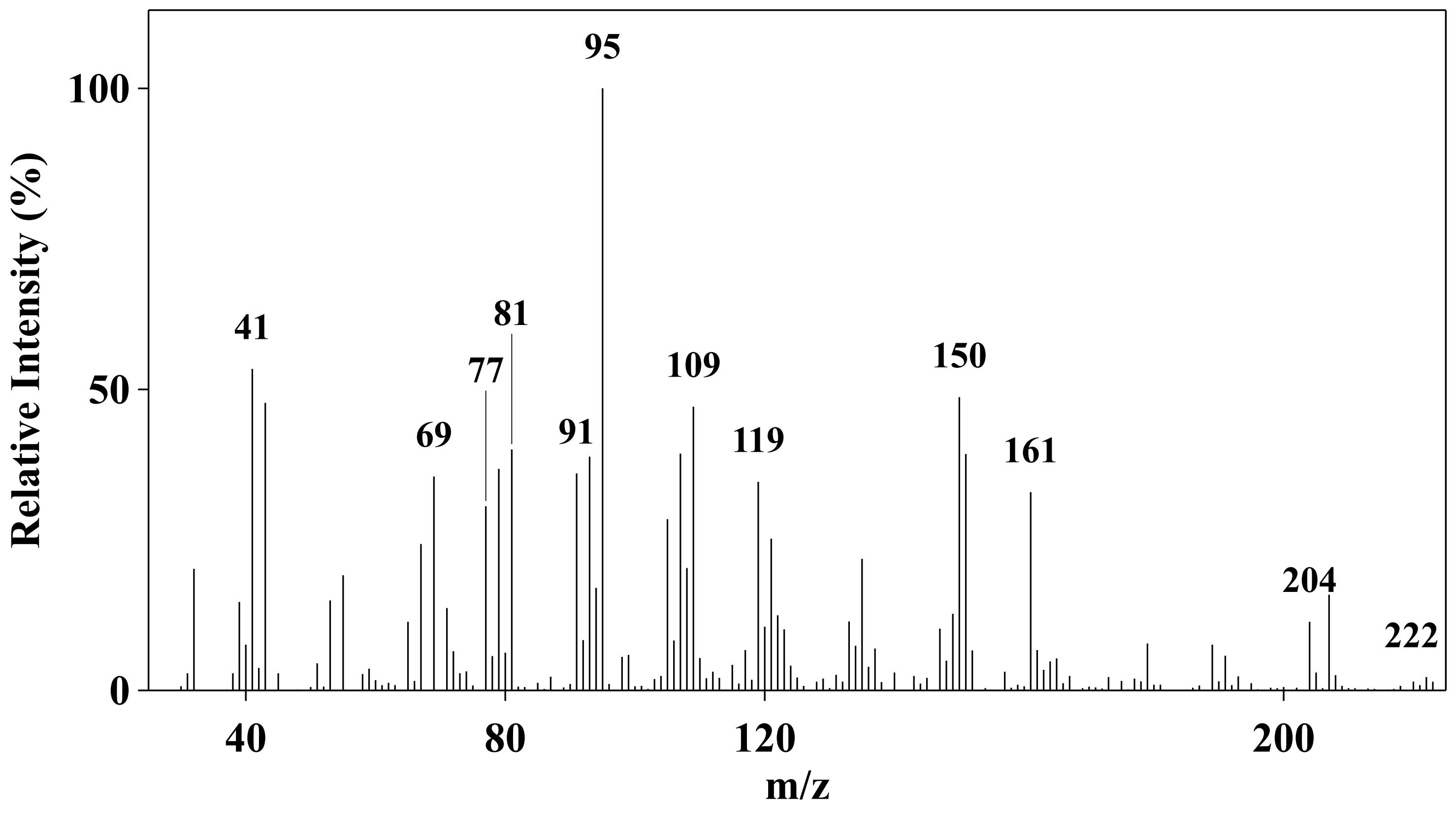


**#26 Copaborneol**

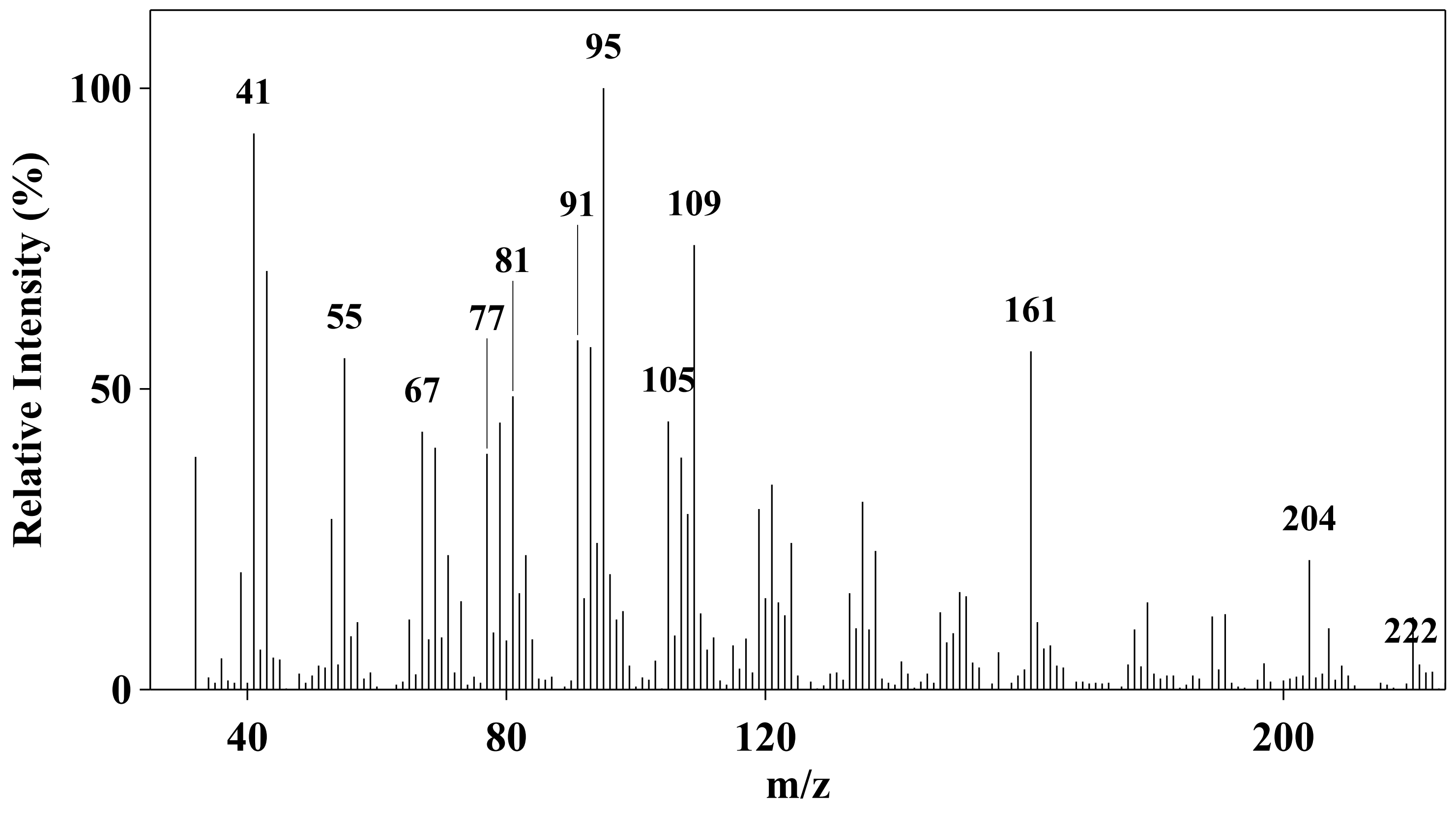


**#27 Dendrobine**

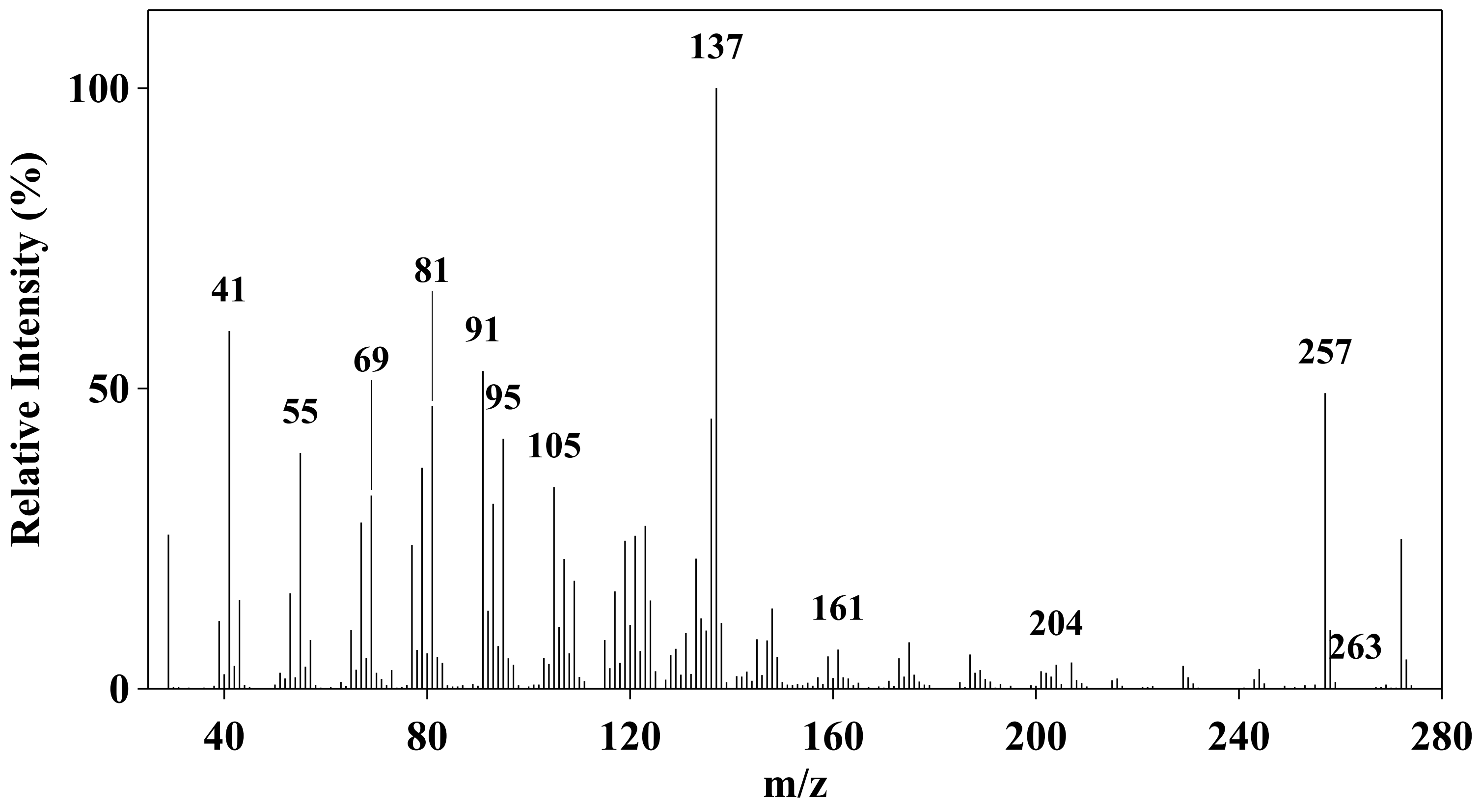
