## Supplementary figures and images for "Functional characterization of terpene synthases from *Dendrobium nobile* provides insights into the biosynthesis of copaborneol, a proposed dendrobine precursor"

### Fig. S2

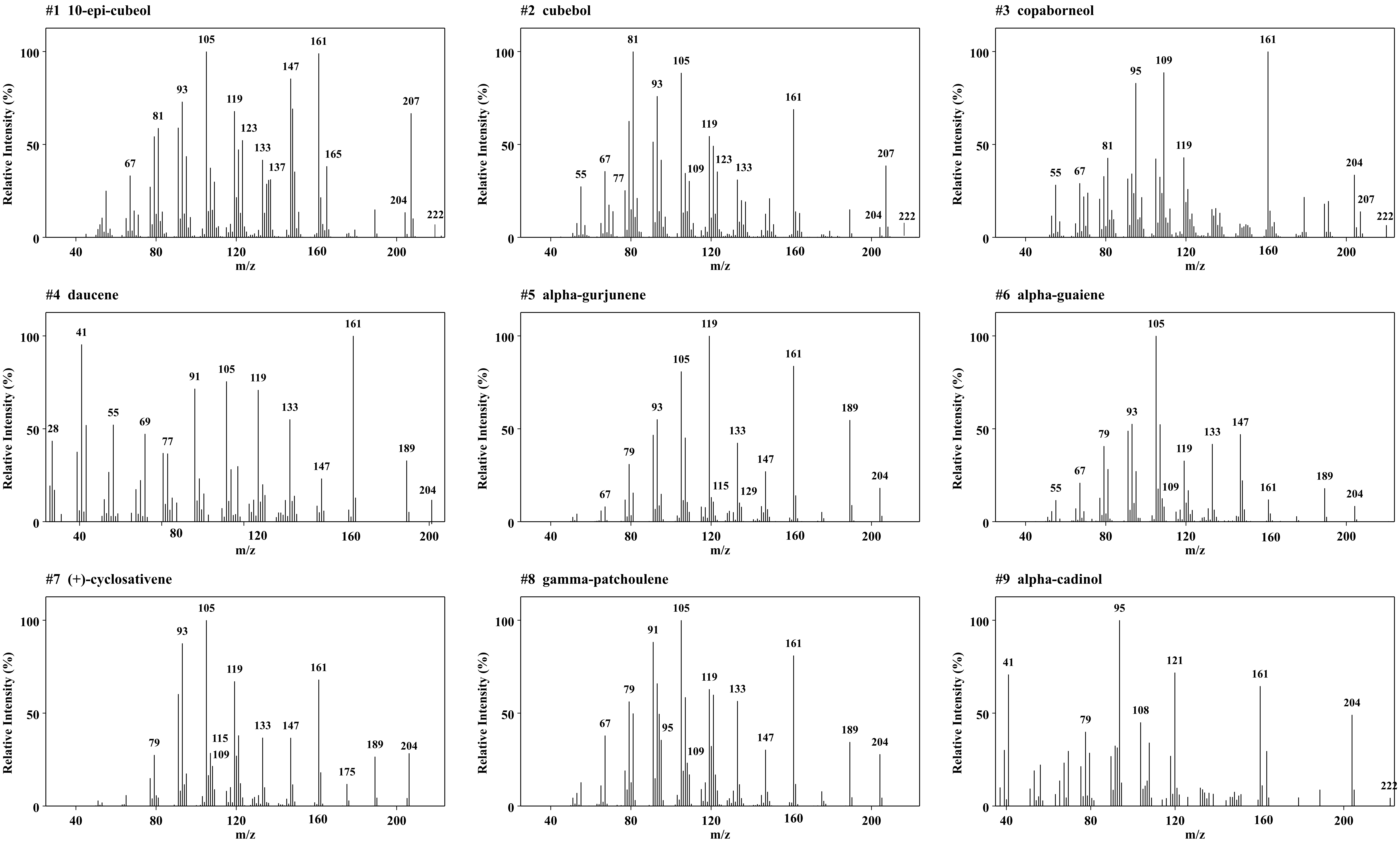

### Fig. S3

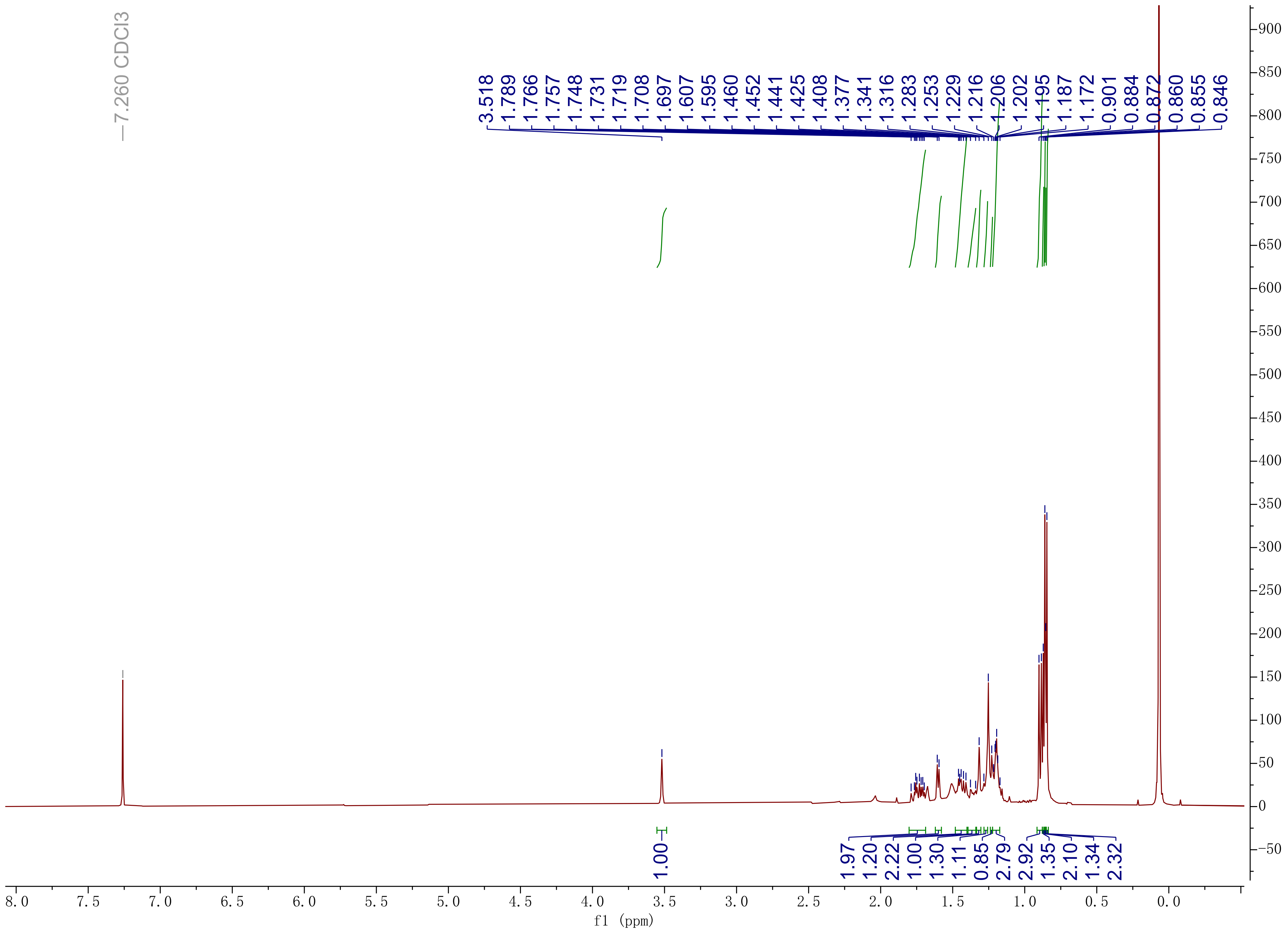

### Fig. S4

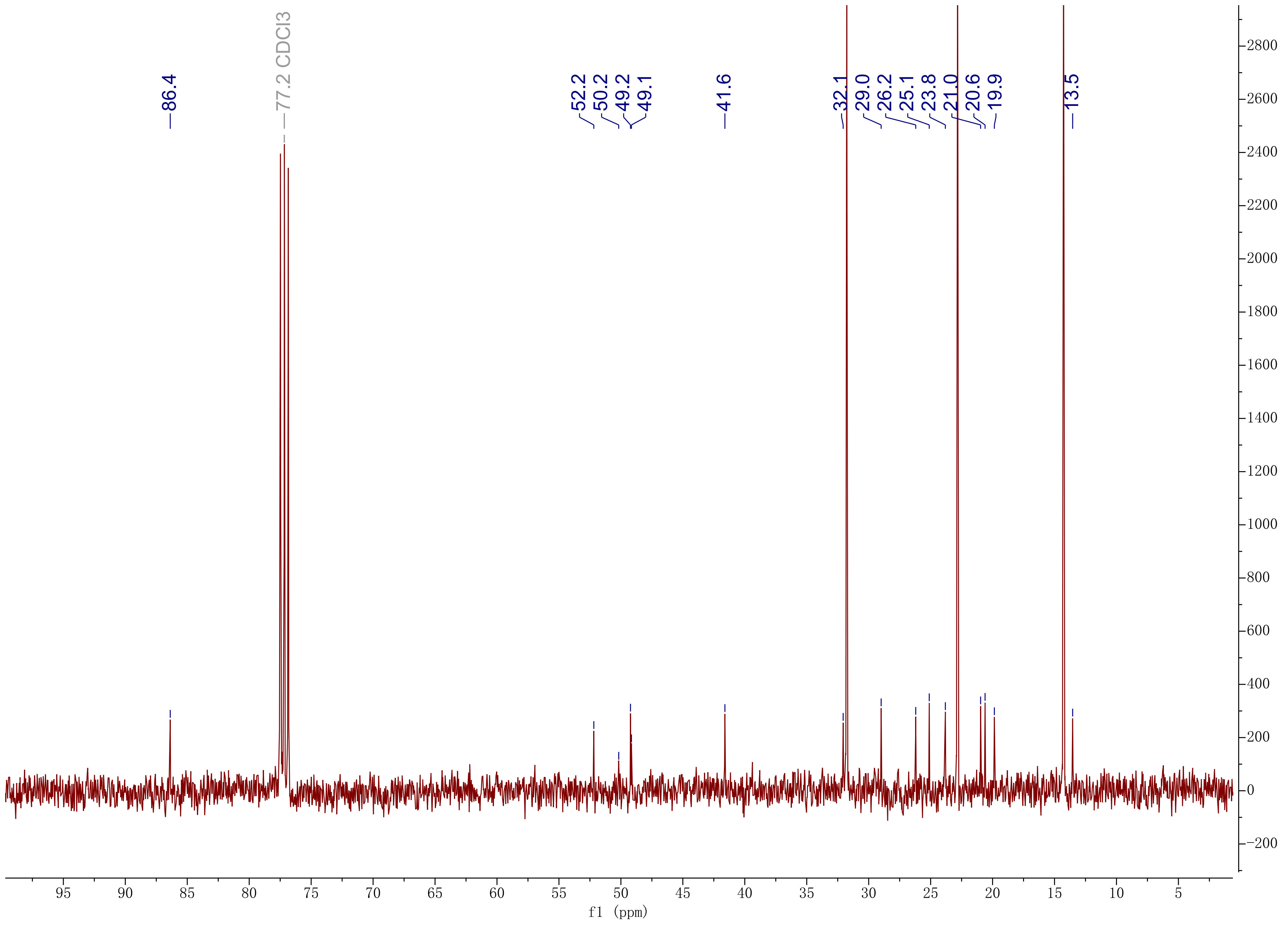
